# Conserved Cdk inhibitors show unique structural responses to tyrosine phosphorylation

**DOI:** 10.1101/2021.02.04.429742

**Authors:** Jacob B. Swadling, Tobias Warnecke, Kyle L. Morris, Alexis R. Barr

## Abstract

Balanced proliferation-quiescence decisions are vital during normal development and in tissue homeostasis and their dysregulation underlies tumorigenesis. Entry into proliferative cycles is driven by Cyclin/Cyclin-dependent kinases (Cdks). Conserved Cdk inhibitors (CKIs), p21^Cip1/Waf1^, p27^Kip1^ and p57^Kip2^, bind to Cyclin/Cdks and inhibit Cdk activity. p27 tyrosine phosphorylation, in response to mitogenic signalling, promotes activation of CyclinD/Cdk4 and CyclinA/Cdk2. Tyrosine phosphorylation is conserved in p21 and p57, although the number of sites differs. We use molecular dynamics simulations to compare the structural changes in Cyclin/Cdk/CKI trimers induced by single and multiple tyrosine phosphorylation in CKIs and their impact on CyclinD/Cdk4 and CyclinA/Cdk2 activity. Despite shared structural features, CKI binding induces distinct structural responses in Cyclin/Cdks and the predicted effects of CKI tyrosine phosphorylation on Cdk activity are not conserved across CKIs. Our analyses suggest how CKIs may have evolved to be sensitive to different inputs to give context-dependent control of Cdk activity.

---

During development and in tissue homeostasis, the controlled release of cells from a quiescent (resting, G0) state into proliferative cycles is vital to achieve and maintain a healthy state. Dysregulation of these pathways underpins abnormal development and tumorigenesis. Cell proliferation can be induced by multiple signals, including mitogenic growth factors, the extracellular matrix and cell-cell contacts. Despite differences in upstream signalling, common to signals that promote proliferation is the downstream activation of Cyclin-dependent kinases (Cdks), that drive entry into and through the cell cycle (G1, S, G2 and mitosis).^1^ How context-dependent signals are translated into changes in Cdk activity is not fully understood.

Cdks are subject to layers of regulation to ensure quiescent cells only enter the cell cycle in response to appropriate environmental cues.^2^ Cdk activation requires Cyclin binding and T-loop phosphorylation in the Cdk-activating segment by Cdk-activating kinase (CAK). Mitogenic signalling promotes Cyclin D expression, which binds Cdk4 (or Cdk6) to initiate progression from G0 into G1. Downstream of CyclinD/Cdk4, Cyclin E then Cyclin A expression activates Cdk2 and promotes the transition of cells from G1 into S-phase (DNA replication). Cdks can be inhibited via Wee1 and Myt1 mediated phosphorylation, which is opposed by Cdc25 phosphatases.^3,4^ Two families of Cdk inhibitors (CKIs) also inhibit Cdk activity: the INK4 and Cip/Kip proteins. Cip/Kip family members (p21^Cip1^, p27^Kip1^ and p57^Kip2^) are intrinsically disordered proteins (IDPs) that fold upon binding to Cyclin/Cdk dimers to inhibit Cdk activity. CDK inhibition is mediated by the Cip/Kip providing a physical block, via the RXL motif, to the substrate-binding site on the Cyclin subunit, and by disrupting the ATP-binding site in the Cdk - either by insertion of the Cip/Kip 3_10_ helix to block ATP binding, in the case of Cdk2,^5,6^ or by displacing the N-lobe of Cdk4 to prevent ATP binding.^7^ Coordination of Cdk-activating and -inhibiting activities is required to prevent unscheduled Cdk activity and promote timely cell cycle entry.

Mitogenic signalling not only stimulates Cyclin D expression but also initiates the removal of p27. Levels of p27 protein are high in quiescent cells and help maintain the quiescent state.^8–10^ p27 binding to CyclinD/Cdk4 releases the activation segment of Cdk4 to permit CAK phosphorylation of CyclinD/Cdk4.^7^ However, the CAK phosphorylated CyclinD/Cdk4/p27 remains inactive. Mitogen signalling activates receptor tyrosine kinases, such as Janus kinase 2 (JAK2) and non-receptor tyrosine kinases (NRTKs), including the proto-oncogenes, Src, Abl, Yes and Lyn. These tyrosine kinases phosphorylate p27 on one or more of three tyrosine residues that reside in the Cdk-binding domain: Y74 (Src, Yes), Y88 (all NRTKs) or Y89 (Abl, Lyn).^11^ For Cdk4, p27^Y89^ phosphorylation promotes activation of Cdk4 by CAK^12^ and Y74 phosphorylation destabilises the interaction between p27 and CyclinD/Cdk4, allowing Cdk4 to become active and initiate cell cycle entry. For Cdk2, phosphorylation of Y88 in p27 leads to the ejection of the 3_10_ helix from the Cdk2 active site, partially activating Cdk2.^13^ This initiates a positive feedback loop where CyclinA/Cdk2 intramolecularly phosphorylates p27^T187^ which promotes p27 recognition, ubiquitination and subsequent degradation by SCF^Skp2^.^14^

The Cip/Kip proteins differ in their transcriptional regulation. Levels of p27 increase in mitogen-starved cells, or cells induced into quiescence by contact inhibition, and are regulated by the transcription factor, FOXO1.^15^ p21 is an early target of p53 and mediates quiescence in response to intrinsic DNA damage^16–20^ and p57 is encoded by an imprinted gene (CDKN1C) and its expression peaks during organogenesis.^21–26^ These differences in expression reflect their unique roles in proliferation control. However, all three Cip/Kips also display overlapping patterns of tissue expression^27^ and, although divergent in their Ctermini, are highly conserved within their N-terminal domains^28,29^ – the region that mediates Cyclin/Cdk binding and that is sensitive to tyrosine phosphorylation. This suggests that all three interact with Cyclin/Cdks in a similar way. Indeed, NRTKs can phosphorylate all three proteins in vitro.^30^

The number and distribution of tyrosine sites in the N-termini of the Cip/Kip proteins differ, with p27 having three sites (Y74, Y88, Y89), p57 two sites (Y63, Y91) and p21 just one (Y77) suggesting that different Cip/Kips might show unique responses to tyrosine phosphorylation. Recent crystal structures comparing p21 and p27 bound to CyclinD/Cdk4 indicate that the lack of a p27^Y74^ equivalent in p21 could be a key difference between how p21 and p27 regulate Cdk4 activity, such that even after NRTK activation, p21 would remain a Cdk inhibitor.^7^ This may explain how p21 is able to induce quiescence after intrinsic DNA damage, even in the presence of continued growth factor signalling.^19,31,32^ Systematic analyses of the interactions between Cyclin/Cdk dimers and the three Cip/Kips, and the role of tyrosine phosphorylation in regulating Cdk4 and Cdk2 activity are lacking, hampering our understanding of how Cdk activity is regulated in different contexts. In addition, in p27 and p57, where multiple tyrosine phosphorylation sites exist, it is not known if a hierarchy of phosphorylation exists or if cooperativity exists between sites.

Classical (cMD) and accelerated molecular dynamics (aMD) simulations have been applied to investigate the folding and binding of p27 in solution^33^ and revealed how p27 exhibits intrinsic flexibility such that buried tyrosine residues can be phosphorylated, promoting the ejection of the 3_10_ helix from the Cdk2 active site, initiating CyclinA/Cdk2 activation.^34–36^ Whether similar flexibility exists in p57 and p21 is unknown. Here, we have used fully atomistic simulations to systematically compare the binding of p21, p27 and p57 to CyclinA/Cdk2 and CyclinD/Cdk4 complexes and to probe the structural changes mediated by single and multi-site tyrosine phosphorylation across all three Cip/Kips in the regulation of Cdk activity. Despite seemingly conserved mechanisms of Cip/Kip binding to Cyclin/Cdk dimers, we show that p21 binding to CyclinA/Cdk2 has a more significant effect on the disruption of hydrogen bonding at the Cyclin/Cdk interface, suggesting an additional mechanism by which p21 could influence Cdk2. Our simulations reveal that tyrosine phosphorylation of the Cip/Kips induces non-conserved, non-intuitive structural changes, confirming that it is insufficient to extrapolate from what we know regarding p27 phosphorylation to p21 and p57 function. We also show how Cip/Kip binding to CyclinA/Cdk2 can block phosphorylation by Wee1 and Myt1 kinases, imposing an order of events on Cdk2 regulation. Together, our analyses show how p21, p27 and p57 have evolved distinct structural responses to tyrosine phosphorylation that would enable cells to respond specifically to context-dependent signalling.

## Methods

### Model construction

The starting structure of p27/Cdk2/CyclinA was taken from the crystal structure (1JSU).^5^ The derived p21 and p57 bound ternary complexes were constructed using the Phyre2 structure prediction tool and templated on to the 1JSU crystal structure.^37^ Models were constructed of doubly protonated phosphorylated tyrosine in each model - for p27, Y88/Y89/Y74 were phosphorylated and each combination therein. For p57 Y91/Y63, and for p21 Y77 were phosphorylated. All models were parameterised using the Amber FF14SB potentials for canonical proteins.^38^ Missing potentials for the phosphorylated tyrosine residues was provided by Homeyer *et al.*^39^ Models were solvated with 14Å of TIP3P water and neutralised with NaCl. Cdk4/CyclinD/CKI models were similarly built from the crystal structure (6P8E).^7^ As the 3_10_ terminal region of p27 was not crystallised in complex with Cdk4/CyclinD, this was added to make a direct comparison with the Cdk2/CyclinA models.

### Classical molecular dynamics

Energy minimisation was performed for 2000 steps using combined steepest descent and conjugate gradient methods. Following minimisation, 20 ps of classical molecular dynamics (cMD) was performed in the NVT ensemble using a Langevin thermostat^40^ to regulate the temperature as we heated up from 0 to 300 K. Following the heat-up phase, we performed 500 ns of cMD in the isobaric/isothermal (NPT) ensemble using the Berendsen barostat^41^ to maintain constant pressure. All simulations were performed using GPU (CUDA) Version 18.0.0 of PMEMD^42–44^ with long-range electrostatic forces treated with Particle-Mesh Ewald summation.^45^

### Accelerated molecular dynamics

Using information gathered from the cMD simulations regarding the time-averaged potential energy of the system and dihedral energy, we performed accelerated molecular dynamics (aMD)^46^ on all models. The aMD method was employed to identify transitions to conformational states that might otherwise be inaccessible within the time-scale used in cMD. aMD can capture high energy transition states by modifying the potential energy surface through the addition of a non-negative boost potential ∆*V* (*r*) to the original potential *V* (*r*), whenever *V* (*r*) is below a pre-defined energy level. The modified potential is related to the true potential, bias potential and boost energy by:

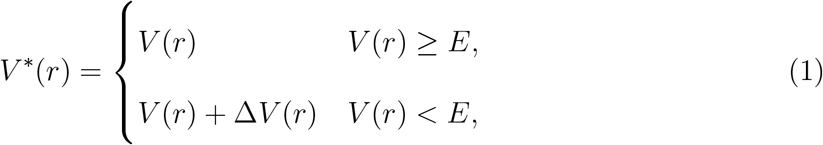

The choice of ∆*V* (*r*) is given by:

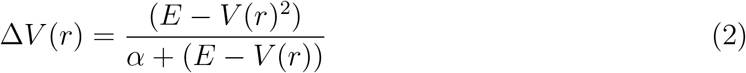

where *α* is a tuning parameter that determines how deep the modified potential energy basin is. *E* and *α* were selected using the following equations:

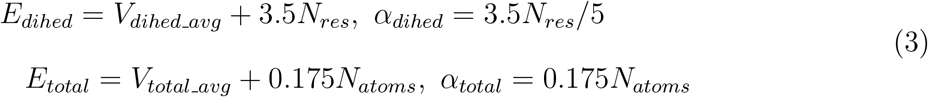

The correct canonical ensemble average of the system was obtained by re-weighting using cumulant expansion to the second order which has been demonstrated to recover the most accurate free energy profiles within statistical errors of ~ *k_B_T*, particularly when the distribution of the boost potential exhibits low anharmonicity (*i.e.*, near-Gaussian distribution), such as the case here.^47^ The Canonical ensemble distribution *p*(*A*), can be recovered using:

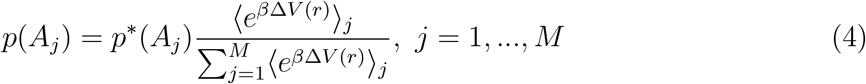

where *M* is the number of bins, *β* = 1*/k_B_T* and (*e^β^*^∆*V*^ ^(*r*)^)_*j*_ is the ensemble averaged Boltzmann factor of ∆*V* (*r*) for simulation frames found in the *j^th^* bin. The ensemble-averaged reweighting factor can be approximated using cumulant expansion to the second order:^47^

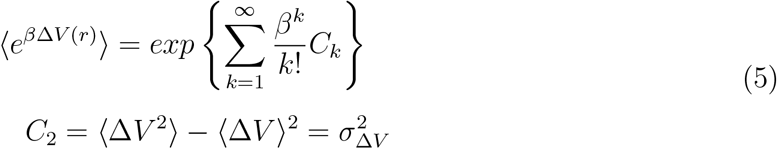

We simulated a total of 20 molecular systems using cMD and aMD, totalling over 20*µs* of simulation, of which the details are given in Supplementary Table 1. Y2P denotes a twice protonated phosphorylation of tyrosine, and T2P denotes a twice protonated phosphorylation of threonine. e.g. Y2P88 indicates the tyrosine residue at residue number 88 has been phosphorylated with two protonation sites (total charge of −2), as per the nomenclature in.^39^

### Identification of Cip/Kip family members and phylogenetic reconstruction

We searched OrthoDB v10 (https://www.orthodb.org) for p21, p27, and p57 based on their corresponding Uniprot identifiers (P38936, P46527, and P49918, respectively) and downloaded the ortholog group that contained all three proteins (1595421) at the Eukaryota level (1595421at2759). Orthologs were then aligned using MAFFT (mafft-linsi). The resulting protein-level alignment was used to compute a maximum likelihood phylogeny using RAxML (v.8.1.16 -f a -m PROTCATAUTO).

## Results

### Hydrogen bonding networks reveal differences in intermolecular contacts between CyclinA, Cdk2 and CKIs

We first sought to understand the effects on the CyclinA/Cdk2 dimer on binding of the different Cip/Kip proteins to ascertain the similarities and differences between these interactions. The structure of CyclinA/Cdk2 bound to p27 has been solved,^5^ but not for p21 or p57. To understand the key contacts made between Cyclin, Cdk and the inhibitor in the ternary complex, we derived homology models for CyclinA/Cdk2/p21 and CyclinA/Cdk2/p57 (Figure 1). We performed cMD simulations and calculated all intermolecular hydrogen bonds over the final 400 ns of trajectories. Contacts which had a life-time fraction above 0.5 were recorded (Supplementary Tables 2-5) and are superimposed upon the final frame of each trajectory (Figure 2). The selected residues represent the major contacts between the three molecules in the ternary complex, which we compare to the CyclinA/Cdk2 dimer. The hydrogen bonding network reveals three significant areas of interaction between the three molecules: i) between Cdk2 and CyclinA, mediated by hydrogen bonding between Cdk2 R150 and CyclinA E268/E269, ii) between Cdk2 and CKI, predominantly through interactions between Cdk2 E81, L20 and V18 and the CKI, and iii) between CyclinA and CKI, via CyclinA Q254 and W217.

**Figure 1:**
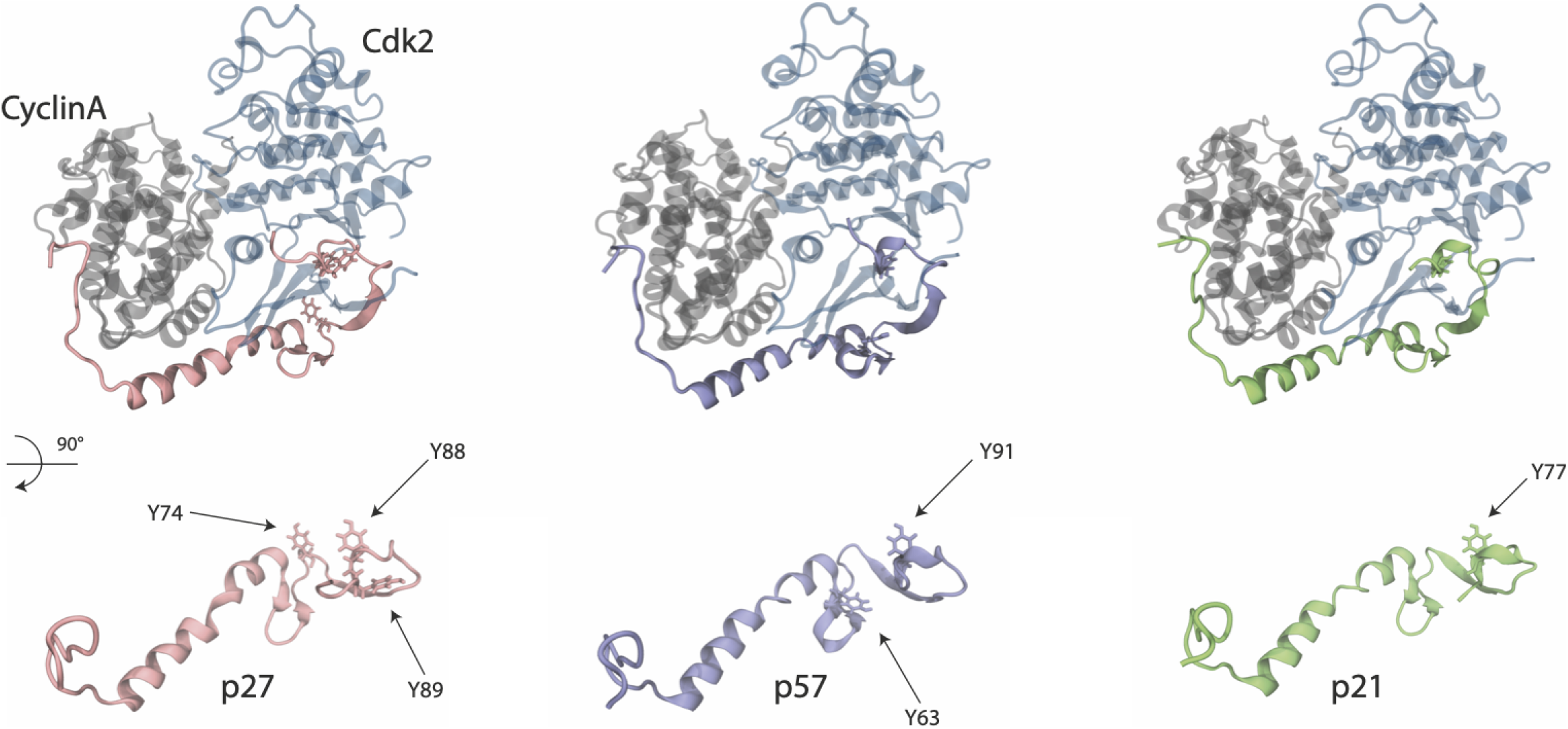
Starting structures of Cdk2 (blue), Cyclin A (grey) in complex with p27 (pink), p57 (purple) and p21 (green) shown as a secondary structure representation. In the lower half of the figure, the three proteins are rotated and tyrosine residues are depicted in a liquorice representation with residue numbers highlighted.

**Figure 2:**
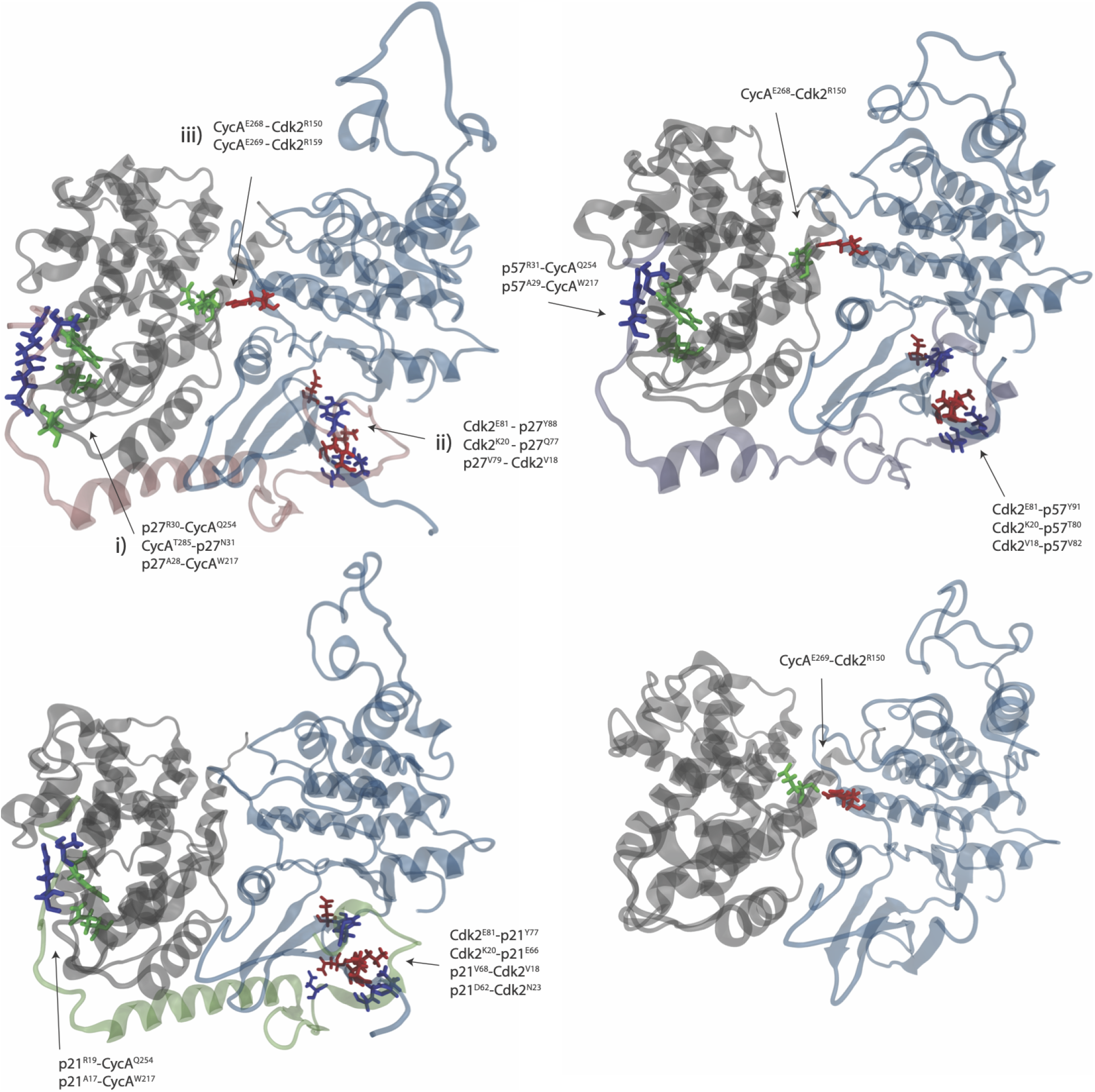
Structures taken from the final frame of each cMD simulation, at 500 ns. Intermolecular hydrogen bonds were calculated over the last 400 ns of simulation and contacts between molecules with a fraction lifetime above 0.5 are displayed on the full structure in the order of hydrogen bond acceptor first and donor second. This simplifies the major, longlived intermolecular contacts between Cdk2, CyclinA and inhibitor. Residues are coloured according to the molecular fragment they belong to: blue is CKI, green is CyclinA, red is Cdk2.

The hydrogen-bonding network revealed by our simulations echoes that observed in the CyclinA/Cdk2/p27 crystal structure.^5^ The simulation of CyclinA/Cdk2/p27 correctly identifies the interaction of the rigid end coil of p27 with the shallow CyclinA groove through the p27 RXL motif and adjacent residues in p27 (R30, N31, A28) with CyclinA Q254, T285 & W217, as shown in Figure 2 (also see Figure 3 in reference^5^ for crystal contacts). Our trajectories also corroborate the importance of binding between p27 Y88 and Cdk2 E81 in the ATP binding site, reported by Russo *et al.*,^5^ where we report this contact to be the single strongest contact made in the trimer (see Supplementary Tables 2-5). Residue R150 in Cdk2 is in the N-terminal region of the T-loop and forms hydrogen bonds to T160, according to.^5^ We observe the hydrogen bond between Cdk2 R150 and CyclinA E268 to be the dominant interaction in the dimer when p27 is bound. We observe changes in H-bonding at the CyclinA/Cdk2 interface that p27 binding induces. Intermolecular contacts between Cdk2 and CyclinA appear strengthened when p27 is bound, through two glutamic acid-arginine residue pairs: E268-R150 and E269-R159.

**Figure 3:**
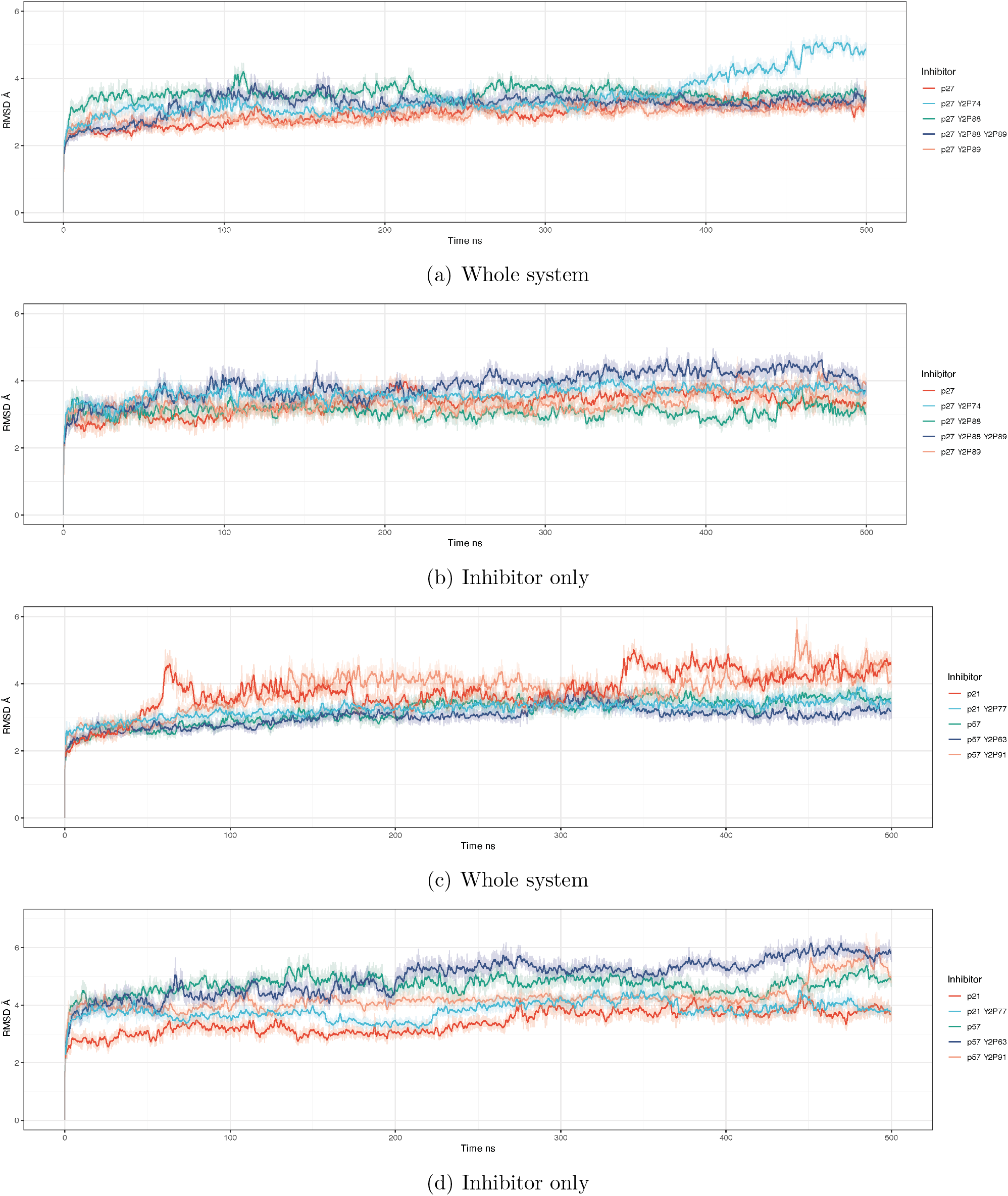
Root mean-squared deviation relative to the starting structure of (a) all Cdk2/CyclinA/p27 models and (b) the isolated p27 inhibitor from each model. (c) Cdk2/CyclinA with p21/p57 partners and (d) the isolated p21/p57 molecules from each model. All models appear to equilibrate after a conservative 25 ns. All time-averaged data was taken with the initial 25 ns of data dropped.

Despite these three distinct areas of contact persisting in all CKI bound complexes, we observe notable differences in the intermolecular interactions between molecules. When p21 is bound, we observe a reduction in the lifetime of the interactions between CyclinA and Cdk2 (to 0.39), suggesting a weakening of the interaction between Cdk2 and CyclinA when p21 is bound, compared to p27 and p57. This could represent an additional mechanism by which p21 might influence Cdk2 activity. Binding of CKI residues to CyclinA is more pronounced in p27 than the other CKIs. Concerning the binding of each CKI to the Cdk2 active site, p21 appears to have the strongest interaction with the presence of a strong intermolecular bond between p21 D62 and Cdk2 N23.

Our homology modelling and calculation of intermolecular hydrogen bonding in the Cdk2/CyclinA/CKI trimers show conserved inhibition mechanisms across all Cip/Kips, through RXL motif dock-ing and occlusion of the Cdk2 active site. We observe differences in the strength of key contacts made between the inhibitors and the dimer, *i.e.* p27 RXL motif having the strongest interaction with Cyclin A and p21 having the strongest interaction with the Cdk2 active site. These differences may be significant in conferring specificity in CKI inhibition and how individual CKIs respond to upstream signalling.

### Tyrosine phosphorylation of conserved Cip/Kips induces distinct conformational changes in the CyclinA/Cdk2/CKI complex

Tyrosine phosphorylation of p27 leads to conformational changes in both the bound Cyclin/Cdk complex and in p27 itself.^35^ To investigate the conservation of tyrosine phosphorylation across Cip/Kip inhibitors, we took an evolutionary approach to determine the gain and loss of these key signalling hubs. p21, p27, and p57 bear distinct signatures of confirmed and putative phosphorylation sites. Focusing on tyrosine residues known to be important for p27 function (Y74, Y88 and Y89) we note that only Y88 is almost universally conserved across vertebrate Cip/Kip family members (Supplementary Figure 1a). The only notable exceptions are p21 orthologs in non-mammalian tetrapods, which all lack Y88 (Supplementary Figure 1b). However, Y88 remains present in p20 (CDKN1D), perhaps suggesting that mammalian p21 functionality might be partitioned between p21 and p20 in these species (Supplementary Figure 1a). Y74 is generally present in p27 orthologs, and the single Cip/Kip copy in non-vertebrates, and is particularly well-conserved in mammals, but largely absent from p21 and p57. Finally, Y89 is restricted to eutherian mammals, *i.e.* absent from marsupials and monotremes and non-mammalian vertebrates.

Each paralogous group also contains several, often well-conserved, residues that could in principle be subject to dynamic phosphorylation (Supplementary Figure 1b). We restricted our analysis to S/T-P sites, which can be phosphorylated by Cdks, MAP Kinases, and GSK3B, all kinases implicated in mitogenic signalling. Some focal residues (e.g. Y74 in p21 and p57) may be compensated by the presence of other residues that are not in an orthologous position but close by, such as Y63 in p57 and T57 in p21, each of which is highly conserved (Supplementary Figure 1b). Indeed, whilst not conserved at the sequence level, Y63 in p57 binds to the Cyclin/Cdk complex in the same position as Y74 in p27, and may be expected to play a similar role (Figure 1). For this reason, here we consider p57^Y63^ as analogous to p27^Y74^. Moreover, while not a tyrosine residue and so not explicitly considered in our analyses, ERK, JNK and GSK3 phosphorylation of p21^T57^ decreases p21 stability and promotes its degradation (shown in Supplementary Figure 2).^48–52^

Having established the conservation of key tyrosine residues in Cip/Kip inhibitors, we wanted to investigate the effect that phosphorylation of these sites would have on the structure of CyclinA/Cdk2/CKI trimers and, ultimately, Cdk2 activity. A systematic investigation into how phosphorylation at each of the three tyrosine residues in p27 alone and in combination could affect these conformational changes and thus if a potential p27 signalling hierarchy or “code” could exist, has yet to be performed. Similarly, whether conformational changes occur in p21 and p57 complexes when conserved tyrosines are phosphorylated remains unknown.

To investigate the dynamics of the different Cip/Kip bound CyclinA/Cdk2 complexes and to identify which structural features warrant further investigation we determined the root mean-squared deviation (RMSD) of each model and that of the inhibitor alone (within the context of CyclinA/Cdk2 binding) in each state of tyrosine phosphorylation we have modelled, relative to the starting structure (Figure 3). We show a single trajectory for each model over a 500 ns time course. It appears that from this general collective variable that each of the models has reached equilibrium after 25 ns. To understand the time-averaged properties of the systems, we drop the first 25 ns from each of the three replicate simulations.

When p27^Y74^ is phosphorylated, p27 shows a significant deviation from the equilibrium average just before 400 ns (Figure 3a, (light blue)). As this is not seen in the RMSD of the inhibitor alone (Figure 3b), it appears that this deviation occurs in the CyclinA/Cdk2 dimer as a consequence of p27^Y74^ phosphorylation. By analysing intermolecular hydrogen bonds over 250-500 ns for the p27^Y2P74^ complex, we see the establishment of hydrogen bonds upon Y74 phosphorylation between CyclinA E268, F267 and Cdk2 R157, R50 which could indicate a change in the conformation after 350 ns. We see little deviation in Y88, Y89, or doubly phosphorylated Y88/Y89 p27, from the equilibrium in these classical MD simulations, indicating that these structures exhibit few conformational changes on these timescales (Figure 3a). Phosphorylation of p27^Y88^ is known to drive the ejection of the 3_10_ helix from the Cdk2 active site.^34,36^ However, this is a slow transition and unlikely to be visible in cMD simulations.^36^

As the p21 and p57 sequences were templated on to the crystal structure for p27 it is not unexpected that they deviate slightly more than that of p27 (compare Figures 3 b and d). Unlike what we observed for p27, for p21 we see CyclinA/Cdk2/p21 jumping from one equilibrium state to another, happening at around 60 and 340 ns (Figure 3c).These shifts seem to occur in the CyclinA/Cdk2 dimer, as they are not observed in the p21 inhibitor (Figure 3d). Phosphorylation of p21^Y77^, which sits in the Cdk2 active site, abolishes these shifts between equilibrium states (Figure 3c). Closer inspection of CyclinA/Cdk2 bound to p21 phosphorylated at Y77 shows the addition of long-lasting intermolecular hydrogen bonding between all three molecules, not observed in the other complexes (see Supplementary Figure 3). For p57^Y91^ phosphorylation, which also sits in the Cdk2 active site, we see the opposite effect to that of p21^Y77^ phosphorylation, where tyrosine phosphorylation promotes transitions between equilibrium states, as compared to the unphosphorylated protein (Figure 3c). This deviation appears to occur in p57 (Figure 3d), and in-part represents the ejection of the 3_10_ helix from the Cdk2 active site when p57^Y91^ is phosphorylated.

Unlike p27^Y74^ phosphorylation, which appears to promote a change in the CyclinA/Cdk2 structure, phosphorylation of p57 at the analogous site (p57^Y63^) leads to a deviation in the structure of p57, not the CyclinA/Cdk2 complex (Figure 3c,d). This suggests potentially important differences in how tyrosine phosphorylation may regulate these conserved Cip/Kips.

In summary, although all Cip/Kip proteins share conserved tyrosine residues, our cMD simulations suggest that phosphorylation of these residues has different effects on CyclinA/Cdk2/CKI complexes. Tyrosine phosphorylation of the residues occupying the Cdk2 active site (p27^Y88,Y89^, p21^Y77^ and p57^Y91^) only induces conformational changes in CyclinA/Cdk2/p21 and CyclinA/Cdk2/p57 on these timescales. However, p21^Y77^ phosphorylation reduces shifts between equilibrium states, while p57^Y91^ phosphorylation increases shifts. Phosphorylation of the more N-terminal residues in p27^Y74^ and p57^Y63^ does induce conformational changes, but in distinct parts of the trimer. Our simulations suggest that tyrosine phosphorylation induces non-intuitive changes in Cyclin/Cdk/CKI trimers.

### Hierarchy and cooperativity of tyrosine phosphorylation in Cip/Kips

A question that arises when considering the phosphorylation of protein residues is whether the amino acid is exposed and accessible to kinases. Many residues are not solvent-exposed in the unphosphorylated state which begs the question of how such residues are modified.^34^ Although the main consideration of this study is to understand the structural and dynamical effects of phosphorylation on these complexes once phosphorylated, it is important to consider the potential of these inhibitors to be modified via phosphorylation. This is particularly relevant here as it has been shown that, in an otherwise unmodified p27 protein, Y88 in p27 is buried against the surface of Cdk2 (Supplementary Figure 4). Yet, experiments have shown that Y88 can still be phosphorylated by NRTKs due to intrinsic flexibility in p27.^35^ However, one would predict that more solvent-exposed residues would have an increased probability of being phosphorylated, may be more likely to be phosphorylated first and may induce conformational changes to promote phosphorylation of other residues, potentially imposing a heirarchy in Cip/Kip phosphorylation.^34^

**Figure 4:**
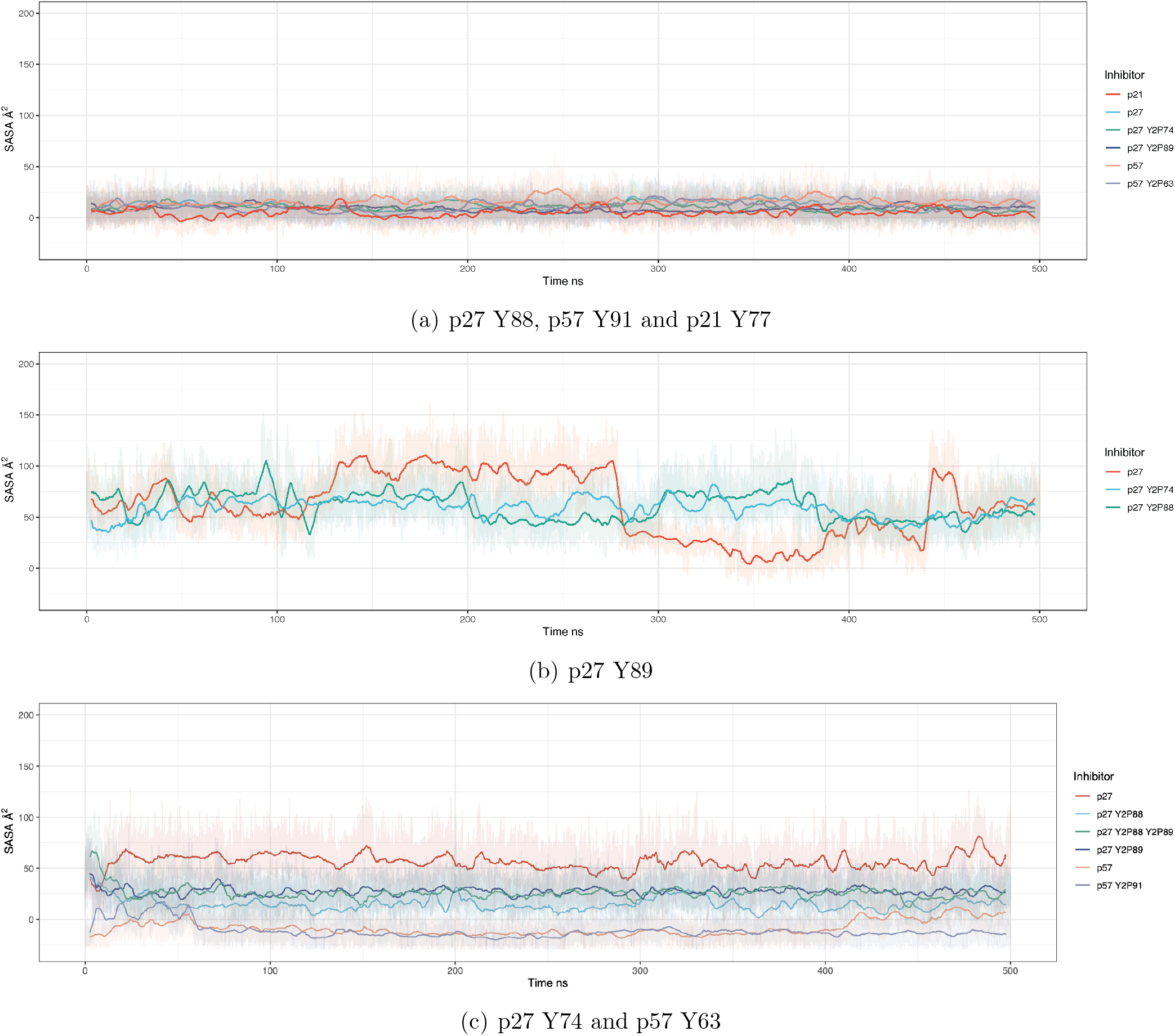
Solvent accessible surface area of three tyrosine residues of interest (a) Y88 in p27, which is positioned within the active site pocket, and the equivalent tyrosine position in p57 and p21 (position Y91 and Y77 respectively). (b) p27 position Y89, positioned within the Cdk2 active site pocket, but only present in p27. (c) Distal from the active site, p27 Y74 and near-equivalent site Y63 in p57.

To determine how solvent-exposed each tyrosine phosphorylation site is in all three Cip/Kips, in the presence and absence of other phosphorylated tyrosines, we calculated the solvent accessible surface area (SASA) using the LCPO algorithm.^53^ From our sequence alignment analysis in Supplementary Figure 1, we have shown the potential importance of several tyrosine residues in terms of their conservation. Previous experimental studies have shown the potential for these tyrosines to be phosphorylated.^54,55^

We first considered the tyrosine residues that sit in the Cdk2 active site. Figure 4a shows the potential for Y88 in p27, and the corresponding positions in p21 and p57, to be phosphorylated, in the presence and absence of phosphorylation of other tyrosine residues (listed on the right-hand side). The SASA of tyrosine at this position (p27^Y88^, p21^Y77^, p57^Y91^) remains close to zero in all models, telling us that throughout these cMD simulations these residues remain solvent inaccessible and buried, and in this state unlikely to be phosphorylated. Figure 4b shows the SASA of Y89, which is only present in p27. Unlike its neighbour Y88, Y89 fluctuates between accessible and inaccessible states in otherwise non-tyrosine phosphorylated p27 (Figure 4b, in red). When Y74 or Y88 are phosphorylated, Y89 remains in a solvent accessible state (see around 280-400 ns in Figure 4b), suggesting cooperativity between these phosphorylation events. Next, we considered the SASA of the more N-terminal tyrosine residues: p27^Y74^ and p57^Y63^. p27^Y74^ remains solvent accessible in its canonical, unmodified, form, whereas p57^Y63^ appears to be buried and inaccessible. Supplementary Figure 4 demonstrates the difference between the accessibility of these two equivalent residues.

To quantify the potential for tyrosine phosphorylation of these residues, we calculated the density of SASA taken from three independent cMD simulations (Supplementary Figure 5). This confirms what we see for residue p27^Y88^, p57^Y91^ and p21^Y77^ - that these residues are buried and inaccessible to solvent, which would imply that they are not available for modification by phosphorylation under these conditions and on these timescales (Supplementary Figure 5a). p27^Y89^ resides in an accessible state in our simulations (Supplementary Figure 5b).

**Figure 5:**
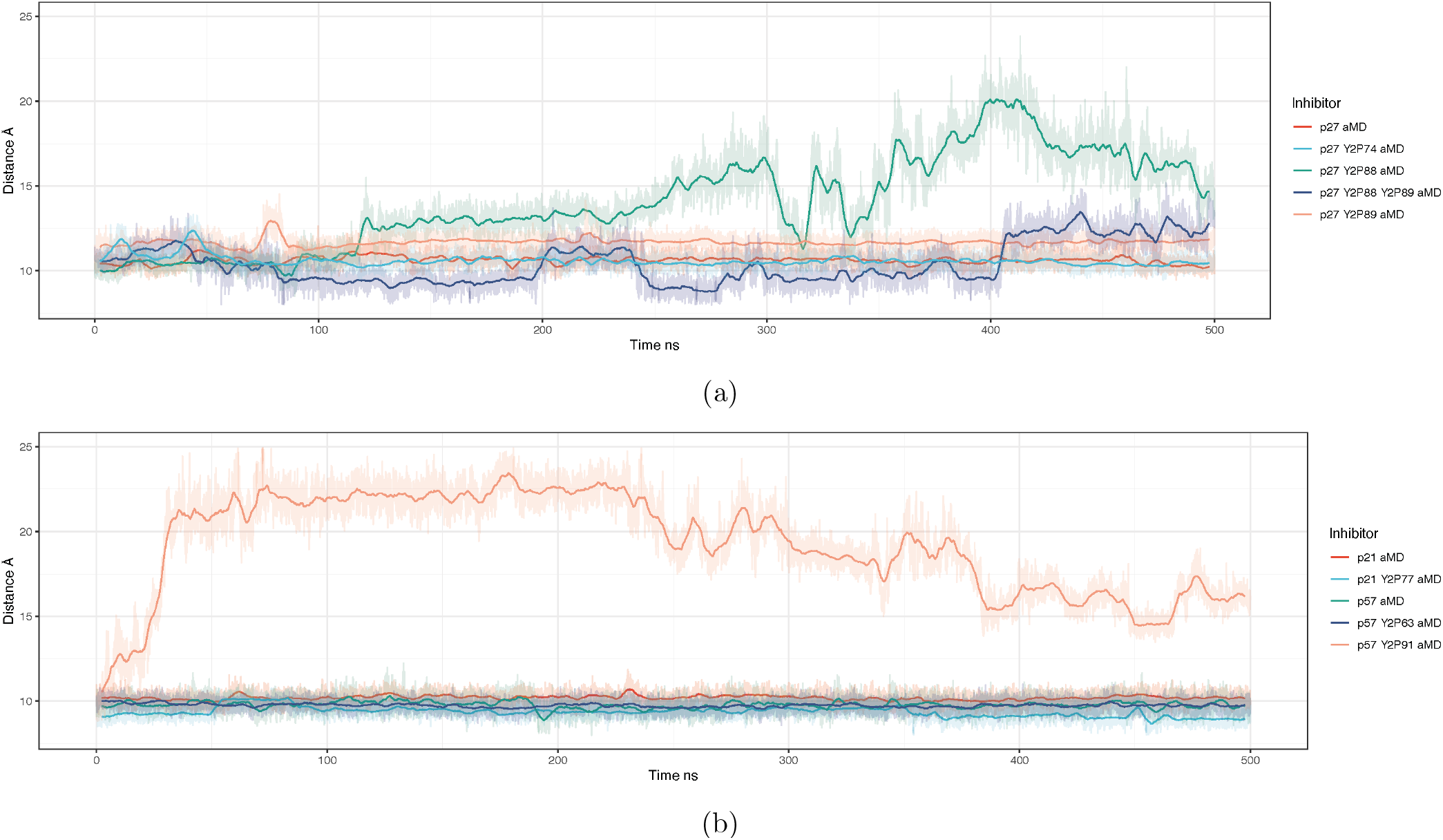
Centre-of-mass distance between the 3_10_ *α*-helices and the active site pocket. Some equilibrium distances appear to be above/below 10Å, because as the phosphorylation state is altered, the centre-of-mass of that group of atoms will be shifted. Of all the various tyrosine phosphorylation states of p27, we only observe dissociation of the inhibitory 3_10_ *α*-helix when Y88 has been phosphorylated. When both Y88 and Y89 are phosphorylated, the *α* helix fluctuates between a bound and partially unbound state, indicating that Y89 might aid in preventing the dissociation of the inhibitor when Y88 is phosphorylated.

Together, these data suggest that the tyrosine residues that sit in the Cdk2 active site (p27^Y88^, p21^Y77^ and p57^Y91^) are inaccessible to phosphorylation. The exception to this is p27^Y89^ which not only remains accessible but also, when phosphorylated, reduces the accessibility of p27^Y74^ for phosphorylation, suggesting cooperativity between phosphorylation of these residues. This mechanism appears to be unique to p27, which is also the only Cip/Kip to have a double YY motif in this position. Since experimental data has shown that tyrosine residues sitting in the Cdk2 active site can be phosphorylated, our simulations indicate that phosphorylation must occur under modifying conditions. For p27^Y88^ that could be brief exposure, or dynamic anticipation, of the residue.^35^ Similar mechanisms could be at play for p21 and p57, or perhaps kinase binding, or other unexplored post-translational modifications, alters residue exposure.

### A predicted route to Cdk2 activation by Cip/Kip release

CyclinA/Cdk2 regains partial activity by ejecting the p27 3_10_ helix from the active site.^35,36^ This event precedes intramolecular phosphorylation of p27 at T187 that targets p27 for proteasomal degradation and would lead to “full” activation of Cdk2.^11^ The 3_10_ helix is present in each of the Cip/Kip inhibitors (Figure 1). To assess the potential for Cdk2 activation in each complex, we monitored the distance between the centre-of-mass of the inhibitor’s 3_10_ helix and the active site pocket. A greater distance suggests a more active kinase, in that the active site pocket is no longer obstructed. Here, we used aMD simulations since it has been previously shown that the 3_10_ helix ejection in p27 happens on slow timescales and is not visible in cMD simulations.^35^ These distances are shown as a time series for each phosphorylation state in Figure 5 from aMD simulations.

Our models suggest that the only circumstances in which the 3_10_ helix blocking the ATP binding site is expelled is in models where the buried tyrosine residue is phosphorylated, *i.e.* positions Y88 in p27 and Y91 in p57. Double phosphorylation of Y88 and Y89 in p27 does not lead to the expulsion of the helix, suggesting that phosphorylation of these adjacent residues have different structural effects (Figure 5). The rate of 3_10_ helix dissociation is greater for p57^Y91^ phosphorylation (Supplementary Figure 6) than for p27^Y88^ phosphorylation. In fact, in p57, when Y91 is phosphorylated, the helix can dissociate during cMD simulations over 500 ns (Supplementary Figure 7). Experimental data shows that, despite being buried, p57^Y91^ can be phosphorylated by Abl kinase, at least *in vitro*.^30^ Therefore, our simulations suggest that whilst there is potentially a low probability of p57^Y91^ phosphorylation, in the absence of other modifying factors, once Y91 is phosphorylated, Cdk2 would be rapidly activated (Figure 5).

**Figure 6:**
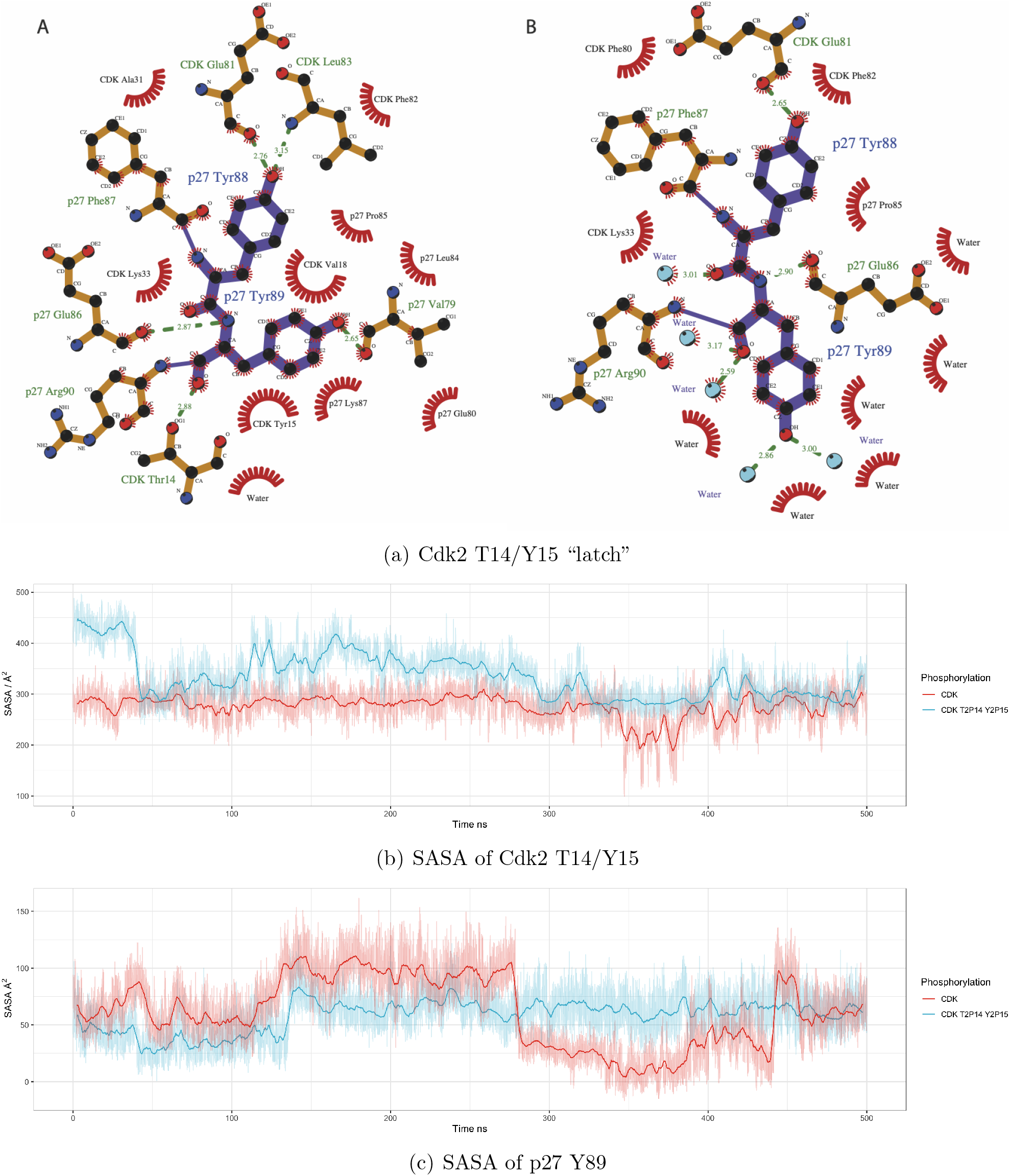
a) 2D rendering of residues Y88 and Y89 in p27 taken from the centroid structure of two conformational states in Fig 4b. b) Distance between p27 residue Y89 and the centre-of-mass of Cdk2 residues T14 & Y15. c) The solvent accessible surface area of p27 Y89, when Cdk2 residue T14 and Y15 are phosphorylated to T2P14 and Y2P15.

**Figure 7:**
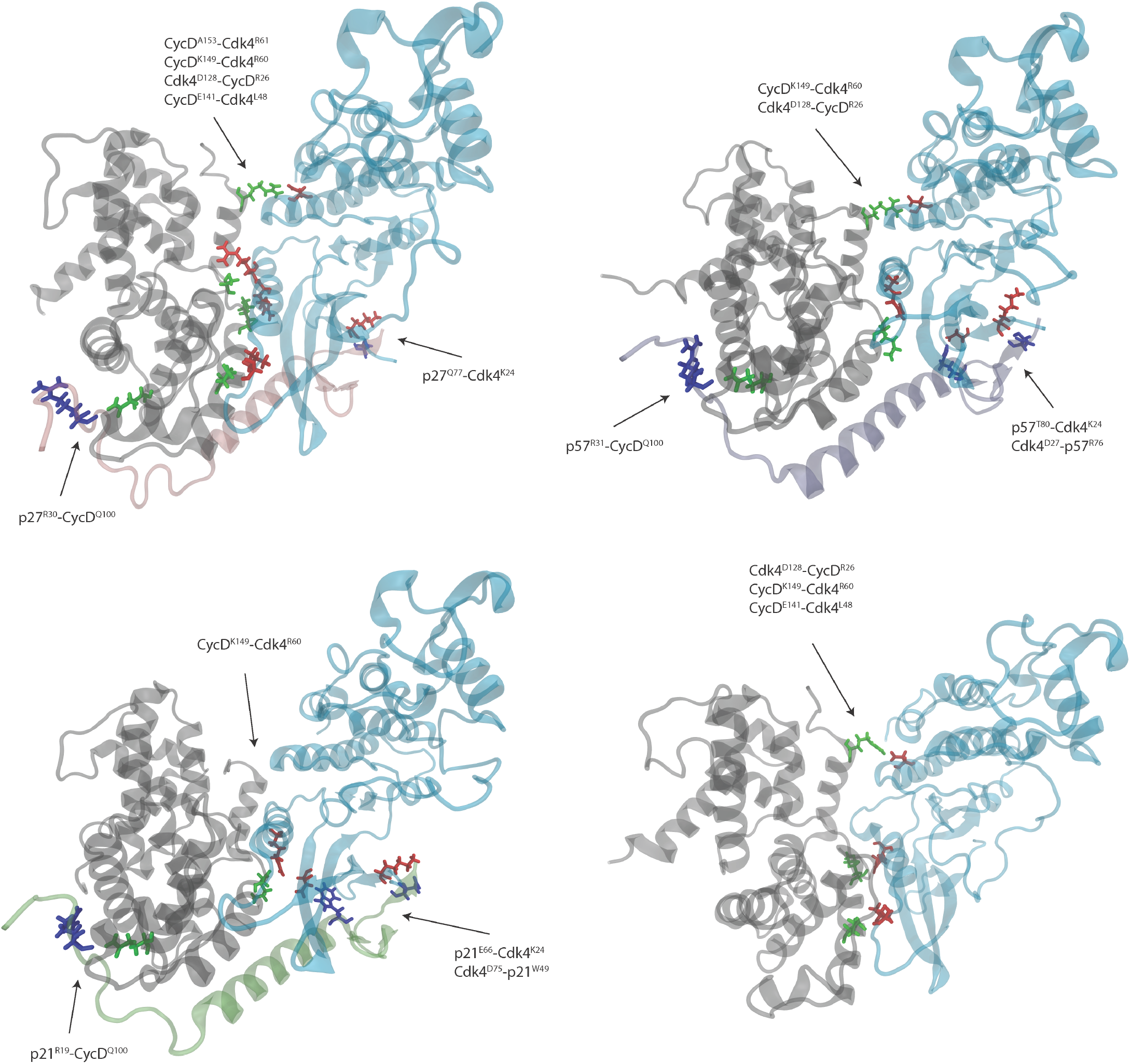
Structures taken from the final frame of each cMD simulation, at 500 ns. Intermolecular hydrogen bonds were calculated over the last 400 ns of simulation and contacts between molecules with a fraction lifetime above 0.5 are displayed on the full structure in the order of hydrogen bond acceptor first, and donor second. This simplifies and shows the major, long-lived intermolecular contacts between Cdk4, CyclinD and their inhibitors. Residues are coloured according to the molecule they belong to: blue is CKI, green is CyclinD, red is Cdk4.

For the case of p21 when the similarly positioned tyrosine Y77 is phosphorylated, we do not observe any dissociation of the helix from the Cdk2 active site (Figure 5). This leads us to believe that there is no reactivation of Cdk2 over these time scales when Y77 in p21 is phosphorylated, as the active site remains obstructed. Also, our previous analyses on the RMSD of p21^Y77^ phosphorylated complexes predicted increased hydrogen bonding within the trimer, suggesting that, even if phosphorylated, p21 may remain a bound inhibitor.

Together, our simulations suggest that the sensitivity to the ejection of the 3_10_ helix from the Cdk2 active site differs between the three Cip/Kip inhibitors with p57 being the most sensitive to release after tyrosine phosphorylation and p21 not being released at all. Moreover, even though Y88 and Y89 are adjacent in p27, their phosphorylation does not have the same effect.

### Inhibition of Cdk2 by Cip/Kips and Wee1 are mutually exclusive

Our models and simulations reveal that p27 binding to CyclinA/Cdk2 blocks access to the inhibitory phosphorylation sites T14 and Y15 in the Cdk2 active-site pocket (Figure 6a). These models, together with published experimental data,^56^ support the idea that p27 has to be removed from CyclinA/Cdk2 before Myt1 and Wee1 can phosphorylate Cdk2. Similarly, if we simulate the phosphorylation of T14 and Y15 in the CyclinA/Cdk2/p27 trimer, the N-terminus of Cdk2 becomes more diffuse, suggesting that it is no longer bound to p27 (Figure 6b).

In Cdk2 that is unphosphorylated at T14 and Y15, the N-terminal domain of Cdk2 is not free and exposed. The red time-series in Figure 6c shows the solvent accessible surface area of p27 Y89. The two states visible, either accessible or not accessible to water, arise due to intermolecular hydrogen bonds between Cdk2 T14 and p27 Y89 that have a lifetime of a couple of hundred nanoseconds. This is shown (Figure 6a) as a two-dimensional LigPlot^57^ of residue positions surrounding p27^Y88,Y89^ taken from the centroid structure from the two conformational ensembles mentioned. When Cdk2 is phosphorylated at residue T14 and Y15, we see this open/close conformational switch removed and the N-terminal domain of Cdk2 becomes unbound to p27 and diffuse (Figure 6c). Together, these data suggest that inhibition of Cdk2 by Cip/Kips and inhibitory kinases are mutually exclusive events.

### Comparison of Cip/Kip binding & release between Cdk2 and Cdk4

The recently solved structures of CyclinD/Cdk4/p27 and CyclinD/Cdk4/p21 allow us to compare Cdk2 and Cdk4 regulation by the Cip/Kip proteins.^7^ We constructed models to compare the binding of all three Cip/Kips to CyclinD/Cdk4 (see Supplementary Figure 8 for a side-by-side comparison of Cdk2-Cdk4 structures). Analysis of RMSD shows CyclinD/Cdk4/CKI models equilibrating after 25 ns of initial simulation, like those we reported for Cdk2/CyclinA/CKI, shown in Supplementary Figure 9. We calculated all intermolecular hydrogen bonds between CyclinD/Cdk4/CKI and observed three main domains of interaction, analogous to those seen in the Cdk2 ternary complex (Figure 7 and Supplementary Tables 6-9), *i.e.* between Cdk4 and CyclinD via the C-helix, between CKI and Cdk4, and between CKI and CyclinD. When compared to CyclinA/Cdk2, the CyclinD/Cdk4 interaction appears to be far stronger, certainly when p27 is bound. Conversely, the CyclinD/CKI interaction is weaker, with a single dominant interaction residue, namely CyclinD^Q100^. Where p27^Y88^, p21^Y77^ and p57^Y91^ binding is the dominant interaction in the Cdk2 models, rep-resenting strong interactions between the 3_10_ helix and the Cdk2 active site, the strongest interacting residues in the Cdk4 models are p27^Q77^, p57^T80^ and p21^E66^ bound to Cdk4^K24^.

**Figure 8:**
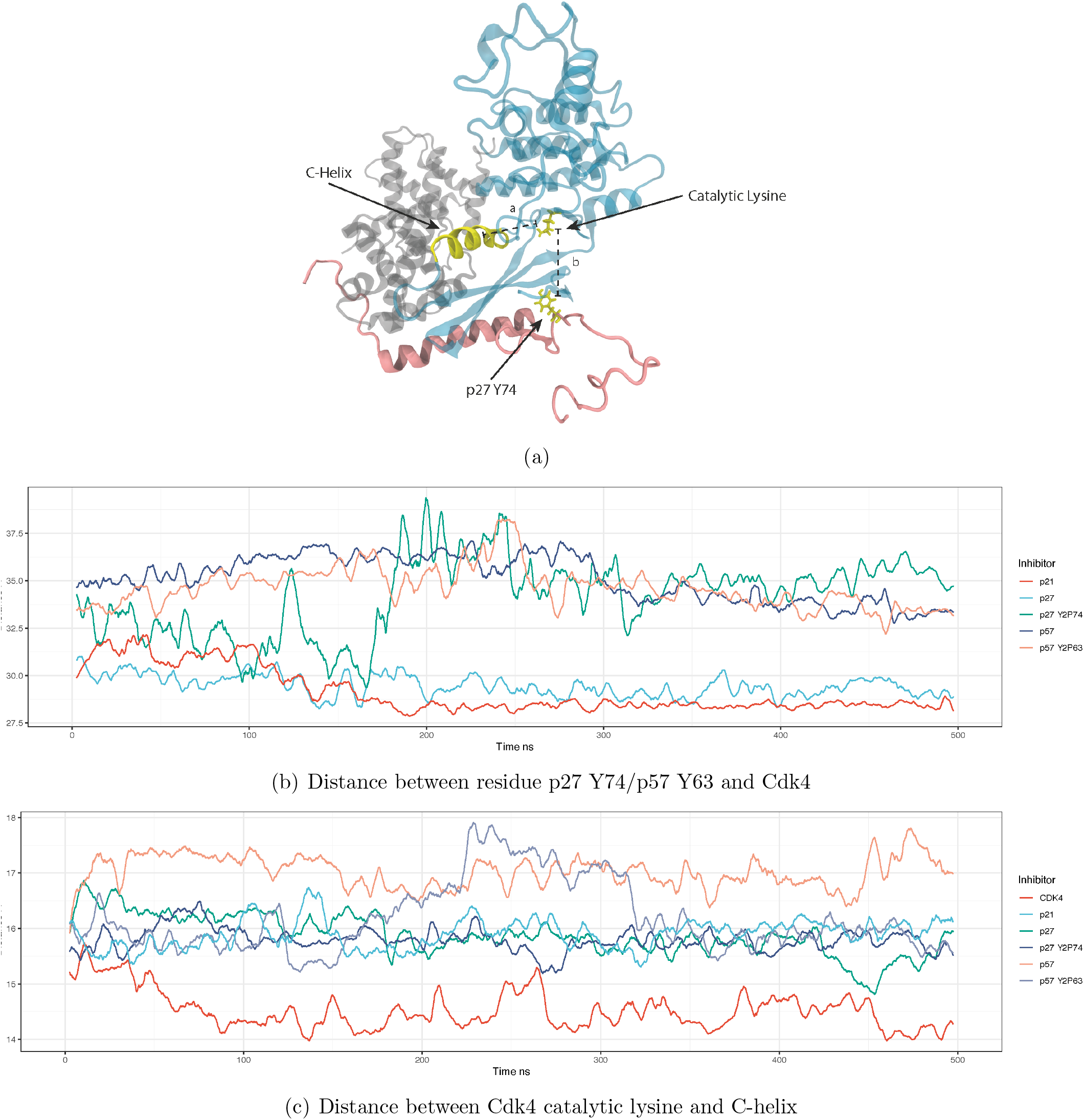
(a) The two distances displayed in yellow on the starting structure of CyclinD/Cdk4/p27, (b) the distance between the equivalent of p27^Y74^ of the three Cip/Kip inhibitors in different states of phosphorylation, and the centre-of-mass of Cdk4. As the dislocation of residue 74 from the complex is known to lead to activation of Cdk4, we show this as an indicator of potential activity. (c) the distance between the catalytic lysine in Cdk4 and the centre-of-mass of the Cdk4 C-helix.

Our analysis of interactions over the course of the cMD simulations reveal differences between the three CKIs when bound to the dimer. p21 and p57 have an interaction close to the Cdk4 active site (p21^W49^ and p57^R76^), which is not observed in p27. It appears that this additional interaction leads to the weakening of the hydrogen bonding between CyclinD/Cdk4 - effectively pulling apart the Cdk4^L48^ - CyclinD^E141^ interaction (Figure 7).

Unlike CyclinA/Cdk2 which is active in the dimeric complex, CyclinD binding alone does not induce catalytic activity in Cdk4.^7^ Compared to CyclinA/Cdk2, where Cip/Kips inhibit kinase activity, Cip/Kip binding can both inhibit and promote CyclinD/Cdk4 activity. Therefore we cannot infer Cdk4 activity from the absence of a Cip/Kip blocking the active site alone - such as the 3_10_ helix in the case Cdk2 inhibition. In Figure 8a, we combine two distances to infer the potential for Cdk4 activity. With both distances, the closer the catalytic lysine is to either the C-helix or the p27^Y74^ (or equivalent) residue, the less active Cdk4 is.

Figure 8b shows the distance between p27^Y74^, the corresponding amino acid in p21 (which is not a tyrosine), p57^Y63^, and the centre-of-mass of Cdk4 (distance b in Figure 8a). This distance tells us whether the tyrosine residue is tightly bound to the Cdk4 molecule, suggesting that the Cdk4 molecule is inactive. Of the models studied, unphosphorylated p21 and p27 are tightly bound at this position suggesting Cdk4 is inactive (Figure 8b). If Y74 is phosphorylated in p27 (green line), Y2P74 transitions from bound to unbound, at around 150 ns and we predict Cdk4 would become more active - consistent with published results.^7^ p57, in both unphosphorylated and Y63 phosphorylated forms, is unbound throughout the simulation, suggesting p57 may be a poor CyclinD/Cdk4 binder.

A second measure of the potential for CyclinD/Cdk4 to be active is the distance between the Cdk4 C-helix and the catalytic lysine residing in the active site (Figure 8c). This is a measure of the “openness” of the active site and a proxy for Cdk4’s ability to accept ATP. Without one of the Cip/Kip family of inhibitors, CyclinD/Cdk4 (red in Figure 8c) exhibits the smallest distance between the C-helix and the catalytic lysine over the course of a 500 ns simulation, suggesting Cdk4 is inactive. Considering both of these distances together, that is the distance between the catalytic lysine and C-helix and between p27^Y74^ and Cdk4 (shown in Figure 8a), allows us to predict Cdk4 activity in these inhibitor-bound states. Taking both measurements into account, we suggest the following order of activity for CyclinD/Cdk4-bound complexes: p21 (inactive) = p27 (inactive) and p27 Y2P74 (partially active) *<* p57 Y2P63 (active) *<* p57 (active).

Our MD simulations are consistent with recent data^7^ showing how phosphorylation of Y74 in p27 bound to CyclinD/Cdk4 can promote activity by inducing conformational changes in the CyclinD/Cdk4 complex. Furthermore, our analyses of p57 bound to CyclinD/Cdk4 suggest that p57 may be a worse inhibitor of CyclinD/Cdk4 activity than of CyclinA/Cdk2 activity.^58,59^

## Discussion

How mitogenic signalling can regulate context-dependent cell proliferation is not fully understood. Here, we have tackled this problem by investigating how mitogens, via the activation of tyrosine kinases, regulate structural changes in Cyclin/Cdk/CKI trimers that ultimately drive Cdk activation and cell proliferation. Our findings reveal that despite shared structural features, tyrosine phosphorylation of CKIs promotes non-conserved and non-intuitive changes in Cyclin/Cdk/CKI complexes, and therefore, in Cdk activity.

Cip/Kip inhibitors are IDPs that fold upon binding to Cyclin/Cdk dimers. Our simulations show that notable differences in the strength of hydrogen bonds between Cyclin/CDK/CKI proteins exist. For example, p21 binding to Cyclin/Cdk appears to destabilise interactions between CyclinA-Cdk2 and CyclinD-Cdk4, which may be an additional mechanism of Cdk inhibition by p21. p21 also has an additional strong hydrogen bond to Cdk2, with N23 in the Cdk2 active site, that is not present in p27 or p57. This, together with our observations, under the conditions simulated here, that phosphorylation of p21^Y77^ in the Cdk2 active site does not promote ejection of the 3_10_ helix of p21, even in aMD simulations, suggests that, under mitogenic signalling, p21 inhibits Cdk2. Together with the fact that p21 also lacks the equivalent p27^Y74^ residue, and so remains a Cdk4 inhibitor on mitogen stimulation, our analyses could explain how p21 is able to maintain cell cycle arrest in the presence of DNA damage and ongoing mitogenic signalling. NMR spectroscopy data suggests that the 3_10_ helix of p21 is released from the Cdk2 active site on Y77 phosphorylation, but that p21 remains a bound inhibitor.^30^ One explanation for the difference between our simulations and experimental measurements could be that the tyrosine kinase (Abl in this case) induces a conformational change in p21 that promotes 3_10_ helix ejection.

Our SASA measurements demonstrate that, with the exception of p27^Y89^, all Cip/Kip tyrosine residues that sit in the Cdk2 active site are inaccessible to phosphorylation. Yet, these sites can be phosphorylated. For p27, previous work demonstrated that p27 exhibits intrinsic flexibility, such that p27 can briefly exist in states where Y88 is exposed for sufficient time to be phosphorylated.^35^ Similar flexibility may exist in p21 and p57. Alternatively, kinase binding may induce conformational changes that promote tyrosine exposure. A third mechanism could be that phosphorylation of other tyrosine sites induces conformational changes that promote exposure of residues on the 3_10_ helix. However, we found no evidence for p27^Y74^-or p57^Y63^-phosphorylation increasing solvent accessibility of p27^Y88^ or p57^Y91^. Conversely, it was recently reported that p27^Y88^-phosphorylation increased the exposure of p27^Y74^,^34^ which we did not observe in our simulations. In the models in^34^ p27^Y89^ was mutated to p27^Y89F^ to mimic previous experimental conditions and therefore this may have affected how p27 behaved in the simulations. Therefore, based on our findings, we favour a combination of intrinsic flexibility and kinase-mediated conformational changes promoting tyrosine exposure.

p27 is the only Cip/Kip to have two adjacent tyrosine residues, Y88 and Y89, in the 3_10_ helix. Despite being neighbours, the two residues behave very differently. Whilst Y88 is buried against Cdk2 and solvent inaccessible, Y89 fluctuates between accessible and inaccessible states, making it more exposed for phosphorylation. However, despite an increased likelihood of phosphorylation, Y89 phosphorylation does not initiate 3_10_ helix ejection under the conditions used in our simulations. In fact, simultaneous phosphorylation of Y89 with Y88 appears to suppress helix ejection and Cdk2 activation. This is intriguing and could be linked to preventing Cdk2 activation and cell proliferation under conditions of high-levels of oncogenic tyrosine kinase signalling.

Our simulations of Cip/Kip inhibitors bound to CyclinD/Cdk4 were able to recapitulate recently reported features of Cdk4 activation and inhibition by p27 and p21.^7^ We were also able to show that p57 binding does not promote as extensive hydrogen bonding between CyclinD and Cdk4 as p27 does, suggesting that p57 may not promote CyclinD/Cdk4 complex assembly in the same way as p27. Our analyses also suggest that p57 may be a poor inhibitor of Cdk4 activity, since in the presence of Y63-phosphorylated or unphosphorylated p57, the distance between the catalytic lysine and the C-helix or p57 is large, suggesting a potentially active Cdk4. Our models suggest that ejection of the 3_10_ helix of p57 from the Cdk2 active site is very sensitive to phosphorylation of p57^Y91^. p57 is expressed at high levels during embryogenesis and its expression decreases, becoming much more restricted at E13.5 and only being expressed in a limited number of adult tissues.^60^ This is unlike p21 and p27 which are more ubiquitously expressed. Together with our data suggesting that p57 may be a poor inhibitor of CyclinD/Cdk4, this suggests that in tissues where both p27 and p57 are expressed, phosphorylation of p27 would be the rate-limiting factor in promoting cell cycle entry.

Recent experimental data has suggested that Cdk4/6 inhibitors being used in the clinic in the treatment of Her2-negative, ER-positive breast cancer, for example Palbociclib, may promote cell cycle arrest by forcing the redistribution of Cip/Kip inhibitors from Cdk4 onto Cdk2.^7^ This uncertainty over the mechanism of action of Cdk4/6 inhibitors stresses the need for an in-depth, mechanistic understanding of how Cdk activity is controlled by Cip/Kip inhibitors, in order to improve patient stratification and motivate the development of improved Cdk inhibitors. Our analyses go some way towards achieving a molecular understanding of Cdk activation required for cell proliferation and provide a basis for further experimental exploration in this area.

## Acknowledgement

This project made use of time on UK Tier 2 JADE granted via the UK High-End Computing Consortium for Biomolecular Simulation, HECBioSim (http://hecbiosim.ac.uk), supported by EPSRC (grant no. EP/R029407/1). JBS is funded by a UKRI Innovation Postdoctoral Fellowship. ARB is supported by MRC funding core-funding MC-A658-5TY60 and a Cancer Research UK CDF: C63833/A25729. TW is supported by MRC core funding grant no. MC-A658-5TY40.

## Supporting Information Available

Accompanying the main article we have included supplementary information on the evolutionary alignment of CKIs and additional molecular dynamics analysis. We have also supplied PDB format starting structures of the molecular models used in this study.

## References

(1) Morgan, D. O. Principles of CDK regulation. Nature 1995, 374, 131.

(2) Pennycook, B. R.; Barr, A. R. Restriction point regulation at the crossroads between quiescence and cell proliferation. FEBS letters 2020, 594, 2046–2060.

(3) Mueller, P. R.; Coleman, T. R.; Kumagai, A.; Dunphy, W. G. Myt1: a membrane-associated inhibitory kinase that phosphorylates Cdc2 on both threonine-14 and tyrosine-15. Science 1995, 270, 86–90.

(4) Nilsson, I.; Hoffmann, I. Progress in cell cycle research; Springer, 2000; pp 107–114.

(5) Russo, A. A.; Jeffrey, P. D.; Patten, A. K.; Massagué, J.; Pavletich, N. P. Crystal structure of the p27Kip1 cyclin-dependent-kinase inibitor bound to the cyclin A–Cdk2 complex. Nature 1996, 382, 325.

(6) Hashimoto, Y.; Kohri, K.; Kaneko, Y.; Morisaki, H.; Kato, T.; Ikeda, K.; Nakanishi, M. Critical role for the 310 helix region of p57Kip2 in cyclin-dependent kinase 2 inhibition and growth suppression. Journal of Biological Chemistry 1998, 273, 16544–16550.

(7) Guiley, K. Z.; Stevenson, J. W.; Lou, K.; Barkovich, K. J.; Kumarasamy, V.; Wijeratne, T. U.; Bunch, K. L.; Tripathi, S.; Knudsen, E. S.; Witkiewicz, A. K. p27 allosterically activates cyclin-dependent kinase 4 and antagonizes palbociclib inhibition. Science 2019, 366.

(8) Nakayama, K.; Ishida, N.; Shirane, M.; Inomata, A.; Inoue, T.; Shishido, N.; Horii, I.; Loh, D. Y.; Nakayama, K.-i. Mice lacking p27Kip1 display increased body size, multiple organ hyperplasia, retinal dysplasia, and pituitary tumors. Cell 1996, 85, 707–720.

(9) Oesterle, E. C.; Chien, W.-M.; Campbell, S.; Nellimarla, P.; Fero, M. L. p27Kip1 is required to maintain proliferative quiescence in the adult cochlea and pituitary. Cell cycle 2011, 10, 1237–1248.

(10) Cheng, T.; Rodrigues, N.; Dombkowski, D.; Stier, S.; Scadden, D. T. Stem cell repopulation efficiency but not pool size is governed by p27 kip1. Nature medicine 2000, 6, 1235–1240.

(11) Jäkel, H.; Peschel, I.; Kunze, C.; Weinl, C.; Hengst, L. Regulation of p27 Kip1 by mitogen-induced tyrosine phosphorylation. Cell Cycle 2012, 11, 1910–1917.

(12) Ray, A.; James, M. K.; Larochelle, S.; Fisher, R. P.; Blain, S. W. p27Kip1 inhibits cyclin D-cyclin-dependent kinase 4 by two independent modes. Molecular and cellular biology 2009, 29, 986–999.

(13) Grimmler, M.; Wang, Y.; Mund, T.; Cilenšek, Z.; Keidel, E.-M.; Waddell, M. B.; Jäkel, H.; Kullmann, M.; Kriwacki, R. W.; Hengst, L. Cdk-inhibitory activity and stability of p27Kip1 are directly regulated by oncogenic tyrosine kinases. Cell 2007, 128, 269–280.

(14) Carrano, A. C.; Eytan, E.; Hershko, A.; Pagano, M. SKP2 is required for ubiquitin-mediated degradation of the CDK inhibitor p27. Nature cell biology 1999, 1, 193–199.

(15) Sakamaki, J.-i.; Daitoku, H.; Yoshimochi, K.; Miwa, M.; Fukamizu, A. Regulation of FOXO1-mediated transcription and cell proliferation by PARP-1. Biochemical and biophysical research communications 2009, 382, 497–502.

(16) El-Deiry, W. S.; Tokino, T.; Velculescu, V. E.; Levy, D. B.; Parsons, R.; Trent, J. M.; Lin, D.; Mercer, W. E.; Kinzler, K. W.; Vogelstein, B. WAF1, a potential mediator of p53 tumor suppression. Cell 1993, 75, 817–825.

(17) El-Deiry, W. S.; Harper, J. W.; O’Connor, P. M.; Velculescu, V. E.; Canman, C. E.; Jackman, J.; Pietenpol, J. A.; Burrell, M.; Hill, D. E.; Wang, Y. WAF1/CIP1 is induced in p53-mediated G1 arrest and apoptosis. Cancer research 1994, 54, 1169–1174.

(18) Heldt, F. S.; Barr, A. R.; Cooper, S.; Bakal, C.; Novák, B. A comprehensive model for the proliferation–quiescence decision in response to endogenous DNA damage in human cells. Proceedings of the National Academy of Sciences 2018, 115, 2532–2537.

(19) Arora, M.; Moser, J.; Phadke, H.; Basha, A. A.; Spencer, S. L. Endogenous replication stress in mother cells leads to quiescence of daughter cells. Cell reports 2017, 19, 1351–1364.

(20) Yang, H. W.; Chung, M.; Kudo, T.; Meyer, T. Competing memories of mitogen and p53 signalling control cell-cycle entry. Nature 2017, 549, 404–408.

(21) Hatada, I.; Mukai, T. Genomic imprinting of p57 KIP2, a cyclin–dependent kinase inhibitor, in mouse. Nature genetics 1995, 11, 204–206.

(22) Matsuoka, S.; Thompson, J. S.; Edwards, M. C.; Bartletta, J.; Grundy, P.; Kalikin, L. M.; Harper, J. W.; Elledge, S. J.; Feinberg, A. P. Imprinting of the gene encoding a human cyclin-dependent kinase inhibitor, p57KIP2, on chromosome 11p15. Proceedings of the National Academy of Sciences 1996, 93, 3026–3030.

(23) Hatada, I.; Inazawa, J.; Abe, T.; Nakayama, M.; Kaneko, Y.; Jinno, Y.; Niikawa, N.; Ohashi, H.; Fukushima, Y.; Iida, K. Genomic imprinting of human p57 KIP2 and its reduced expression in Wilms’ tumors. Human molecular genetics 1996, 5, 783–788.

(24) Yan, Y.; Frisén, J.; Lee, M.-H.; Massagué, J.; Barbacid, M. Ablation of the CDK inhibitor p57Kip2 results in increased apoptosis and delayed differentiation during mouse development. Genes & development 1997, 11, 973–983.

(25) Zhang, P.; Liégeois, N. J.; Wong, C.; Finegold, M.; Hou, H.; Thompson, J. C.; Silverman, A.; Harper, J. W.; DePinho, R. A.; Elledge, S. J. Altered cell differentiation and proliferation in mice lacking p57KIP2 indicates a role in Beckwith–Wiedemann syndrome. Nature 1997, 387, 151–158.

(26) Westbury, J.; Watkins, M.; Ferguson-Smith, A. C.; Smith, J. Dynamic temporal and spatial regulation of the cdk inhibitor p57kip2 during embryo morphogenesis. Mechanisms of development 2001, 109, 83–89.

(27) Ullah, Z.; Lee, C. Y.; DePamphilis, M. L. Cip/Kip cyclin-dependent protein kinase inhibitors and the road to polyploidy. Cell division 2009, 4, 1–15.

(28) Dynlacht, B.; Ngwu, C.; Winston, J.; Swindell, E.; Elledge, S.; Harlow, E.; Harper, J. Methods in enzymology; Elsevier, 1997; Vol. 283; pp 230–244.

(29) Sherr, C. J.; Roberts, J. M. Inhibitors of mammalian G1 cyclin-dependent kinases. Genes & development 1995, 9, 1149–1163.

(30) Huang, Y.; Yoon, M.-K.; Otieno, S.; Lelli, M.; Kriwacki, R. W. The Activity and Stability of the Intrinsically Disordered Cip/Kip Protein Family Are Regulated by Non-Receptor Tyrosine Kinases. Journal of molecular biology 2015, 427, 371–386.

(31) Barr, A. R.; Cooper, S.; Heldt, F. S.; Butera, F.; Stoy, H.; Mansfeld, J.; Novák, B.; Bakal, C. DNA damage during S-phase mediates the proliferation-quiescence decision in the subsequent G1 via p21 expression. Nature communications 2017, 8, 14728.

(32) Yang, X.-P.; Liu, S.-L.; Xu, J.-F.; Cao, S.-G.; Li, Y.; Zhou, Y.-B. Pancreatic stellate cells increase pancreatic cancer cells invasion through the hepatocyte growth factor/cMet/survivin regulated by P53/P21. Experimental cell research 2017, 357, 79–87.

(33) Sivakolundu, S. G.; Bashford, D.; Kriwacki, R. W. Disordered p27Kip1 exhibits intrinsic structure resembling the Cdk2/cyclin A-bound conformation. Journal of molecular biology 2005, 353, 1118–1128.

(34) Henriques, J.; Lindorff-Larsen, K. Protein dynamics enables phosphorylation of buried residues in Cdk2/Cyclin A-bound p27. Biophysical Journal 2020, 119, 2010–2018.

(35) Tsytlonok, M.; Sanabria, H.; Wang, Y.; Felekyan, S.; Hemmen, K.; Phillips, A. H.; Yun, M.-K.; Waddell, M. B.; Park, C.-G.; Vaithiyalingam, S. Dynamic anticipation by Cdk2/Cyclin A-bound p27 mediates signal integration in cell cycle regulation. Nature communications 2019, 10, 1–13.

(36) Rath, S. L.; Senapati, S. Mechanism of p27 unfolding for CDK2 reactivation. Scientific reports 2016, 6, 26450.

(37) Kelley, L. A.; Mezulis, S.; Yates, C. M.; Wass, M. N.; Sternberg, M. J. The Phyre2 web portal for protein modeling, prediction and analysis. Nature protocols 2015, 10, 845.

(38) Maier, J. A.; Martinez, C.; Kasavajhala, K.; Wickstrom, L.; Hauser, K. E.; Simmerling, C. ff14SB: improving the accuracy of protein side chain and backbone parameters from ff99SB. Journal of chemical theory and computation 2015, 11, 3696–3713.

(39) Homeyer, N.; Horn, A. H.; Lanig, H.; Sticht, H. AMBER force-field parameters for phosphorylated amino acids in different protonation states: phosphoserine, phosphothreonine, phosphotyrosine, and phosphohistidine. Journal of molecular modeling 2006, 12, 281–289.

(40) Davidchack, R. L.; Handel, R.; Tretyakov, M. Langevin thermostat for rigid body dynamics. The Journal of chemical physics 2009, 130, 234101.

(41) Berendsen, H. J.; Postma, J. v.; van Gunsteren, W. F.; DiNola, A.; Haak, J. R. Molecular dynamics with coupling to an external bath. The Journal of chemical physics 1984, 81, 3684–3690.

(42) Salomon-Ferrer, R.; Götz, A. W.; Poole, D.; Le Grand, S.; Walker, R. C. Routine microsecond molecular dynamics simulations with AMBER on GPUs. 2. Explicit solvent particle mesh Ewald. Journal of chemical theory and computation 2013, 9, 3878–3888.

(43) Götz, A. W.; Williamson, M. J.; Xu, D.; Poole, D.; Le Grand, S.; Walker, R. C. Routine microsecond molecular dynamics simulations with AMBER on GPUs. 1. Generalized born. Journal of chemical theory and computation 2012, 8, 1542–1555.

(44) Le Grand, S.; Götz, A. W.; Walker, R. C. SPFP: Speed without compromise—A mixed precision model for GPU accelerated molecular dynamics simulations. Computer Physics Communications 2013, 184, 374–380.

(45) Essmann, U.; Perera, L.; Berkowitz, M. L.; Darden, T.; Lee, H.; Pedersen, L. G. A smooth particle mesh Ewald method. The Journal of chemical physics 1995, 103, 8577–8593.

(46) Hamelberg, D.; Mongan, J.; McCammon, J. A. Accelerated molecular dynamics: a promising and efficient simulation method for biomolecules. The Journal of chemical physics 2004, 120, 11919–11929.

(47) Miao, Y.; Sinko, W.; Pierce, L.; Bucher, D.; Walker, R. C.; McCammon, J. A. Improved reweighting of accelerated molecular dynamics simulations for free energy calculation. Journal of chemical theory and computation 2014, 10, 2677–2689.

(48) Foertsch, F.; Teichmann, N.; Kob, R.; Hentschel, J.; Laubscher, U.; Melle, C. S100A11 is involved in the regulation of the stability of cell cycle regulator p21 CIP 1/WAF 1 in human keratinocyte H a C a T cells. The FEBS Journal 2013, 280, 3840–3853.

(49) Colleoni, B.; Paternot, S.; Pita, J. M.; Bisteau, X.; Coulonval, K.; Davis, R. J.; Raspé, E.; Roger, P. P. JNKs function as CDK4-activating kinases by phosphorylating CDK4 and p21. Oncogene 2017, 36, 4349–4361.

(50) Densham, R. M.; O’Neill, E.; Munro, J.; König, I.; Anderson, K.; Kolch, W.; Olson, M. F. MST kinases monitor actin cytoskeletal integrity and signal via c-Jun N-terminal kinase stress-activated kinase to regulate p21Waf1/Cip1 stability. Molecular and cellular biology 2009, 29, 6380–6390.

(51) Hwang, C. Y.; Lee, C.; Kwon, K.-S. Extracellular signal-regulated kinase 2-dependent phosphorylation induces cytoplasmic localization and degradation of p21Cip1. Molecular and cellular biology 2009, 29, 3379–3389.

(52) Rössig, L.; Badorff, C.; Holzmann, Y.; Zeiher, A. M.; Dimmeler, S. Glycogen synthase kinase-3 couples AKT-dependent signaling to the regulation of p21Cip1 degradation. Journal of Biological Chemistry 2002, 277, 9684–9689.

(53) Connolly, M. L. Analytical molecular surface calculation. Journal of applied crystallography 1983, 16, 548–558.

(54) Chu, I.; Sun, J.; Arnaout, A.; Kahn, H.; Hanna, W.; Narod, S.; Sun, P.; Tan, C.-K.; Hengst, L.; Slingerland, J. p27 phosphorylation by Src regulates inhibition of cyclin E-Cdk2. Cell 2007, 128, 281–294.

(55) Hornbeck, P. V.; Zhang, B.; Murray, B.; Kornhauser, J. M.; Latham, V.; Skrzypek, E. PhosphoSitePlus, 2014: mutations, PTMs and recalibrations. Nucleic acids research 2015, 43, D512–D520.

(56) Keaton, M. A.; Bardes, E. S.; Marquitz, A. R.; Freel, C. D.; Zyla, T. R.; Rudolph, J.; Lew, D. J. Differential susceptibility of yeast S and M phase CDK complexes to inhibitory tyrosine phosphorylation. Current Biology 2007, 17, 1181–1189.

(57) Laskowski, R. A.; Swindells, M. B. LigPlot+: multiple ligand–protein interaction diagrams for drug discovery. Journal of Chemical Information and Modeling 2011, 51, 2778–2786.

(58) Lee, M.-H.; Reynisdottir, I.; Massague, J. Cloning of p57KIP2, a cyclin-dependent kinase inhibitor with unique domain structure and tissue distribution. Genes & development 1995, 9, 639–649.

(59) Matsuoka, S.; Edwards, M. C.; Bai, C.; Parker, S.; Zhang, P.; Baldini, A.; Harper, J. W.; Elledge, S. J. p57KIP2, a structurally distinct member of the p21CIP1 Cdk inhibitor family, is a candidate tumor suppressor gene. Genes & development 1995, 9, 650–662.

(60) Creff, J.; Besson, A. Functional versatility of the CDK inhibitor p57Kip2. Frontiers in Cell and Developmental Biology 2020, 8.

