## Supplementary Information for "Conserved Cdk inhibitors show unique structural responses to tyrosine phosphorylation"

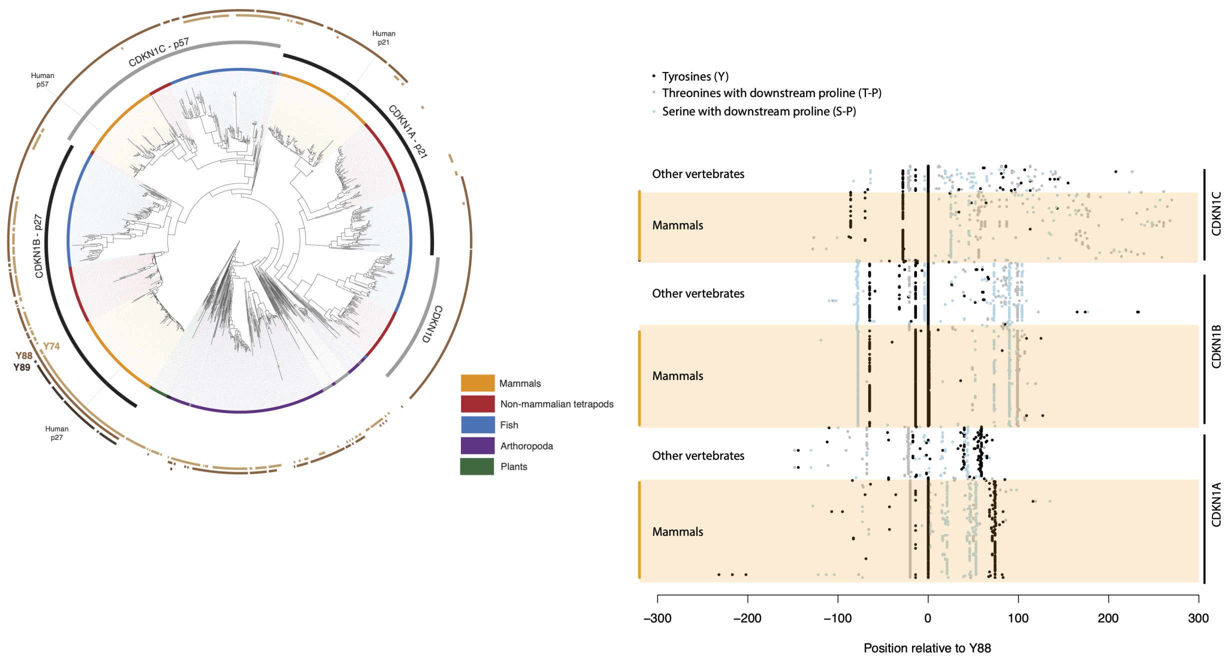


(a) (b)

Supplementary Figure 1: Phylogeny of the Cip/Kip family and conservation of residues subject to phosphorylation. (a) Protein-level maximum likelihood phylogeny of Cip/Kip family members across eukaryotes (see Methods for data provenance). Based both on prior annotation and the phylogenetic splits observed here, CDKN1A (p21), CDKN1B (p27), CDKN1C (p57), and CDKN1D (p20) sequences form coherent monophyletic clades and are labelled accordingly. The presence/absence of key tyrosine residues (Y74, Y88, Y89 in p27) across the tree is indicated. (b) Distribution of potential phosphorylation sites across members of the Cip/Kip family. Tyrosine = black; serine followed by a downstream proline = light blue; threonine followed by a downstream proline = grey. For simplicity, only orthologs previously explicitly annotated as CDKN1A, CDKN1B, or CDKN1C in mammals and non-mammalian tetrapods (birds, reptiles, amphibians) are considered here. Sequences were centred on orthologous position Y88 based on the protein alignment (see Methods). The distance to neighbouring phosphorylation sites was then computed for each sequence independently and therefore faithfully represents the distance along the protein primary sequence rather than along the (gapped) alignment.

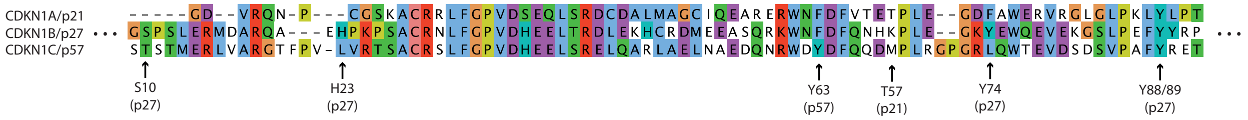


Supplementary Figure 2: Partial alignment of human p21, p27, and p57, extracted from the larger alignment of eukaryotic CIP/KIP orthologs (see main text Methods). Residues of particular interest are highlighted.

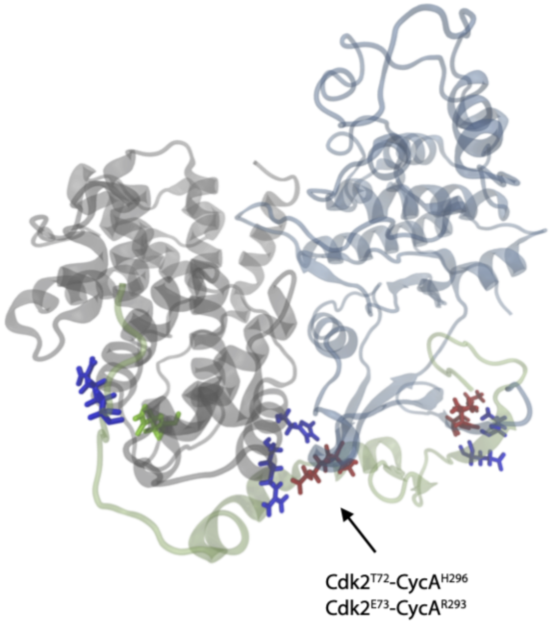


Supplementary Figure 3: CyclinA/Cdk2/p21 structure at 340 ns, where p21 has been phosphorylated at Y77. The structure shows the addition of long-lasting intermolecular hydrogen bonding between all three molecules, not observed in the other complexes (compare to Figure 2).

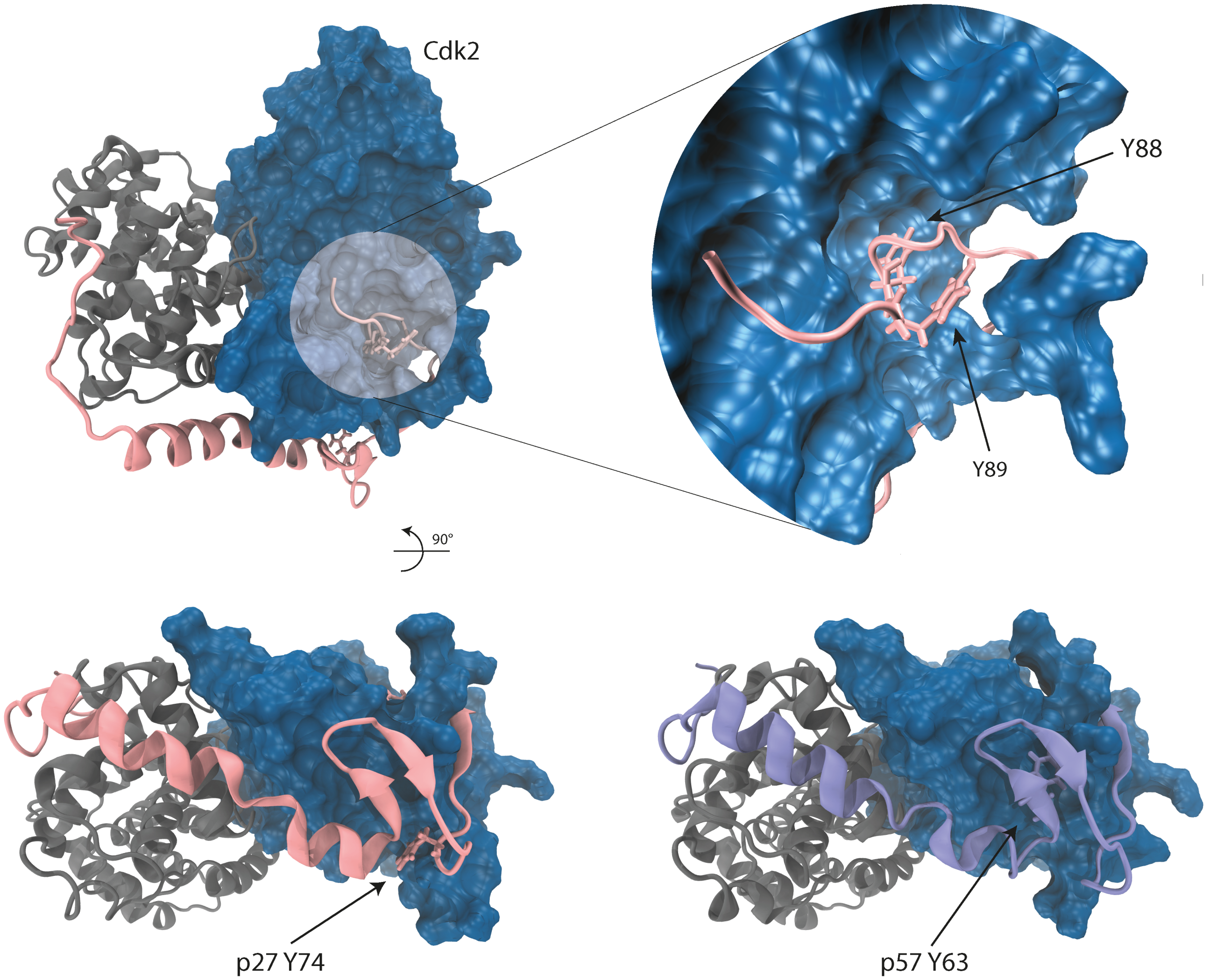


Supplementary Figure 4: Structure of CyclinA/Cdk2/p27 where Cdk2 is displayed as solvent accessible surface area. These structures highlight the inaccessible/buried nature of $p2^{Y88}$, hidden within the Cdk2 active site and obscured by the terminal end of Cdk2. The adjacent Y89 residue is more accessible and potentially more likely to be phosphorylated by a NRTK. The structures for p27 and p57 have been rotated in order to highlight the positions of $p27^{Y74}$ and $p57^{Y63}$.

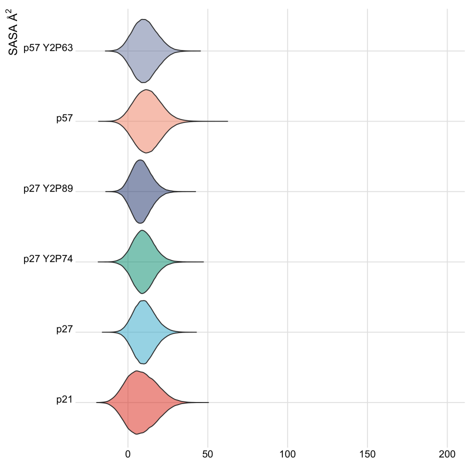


(a)

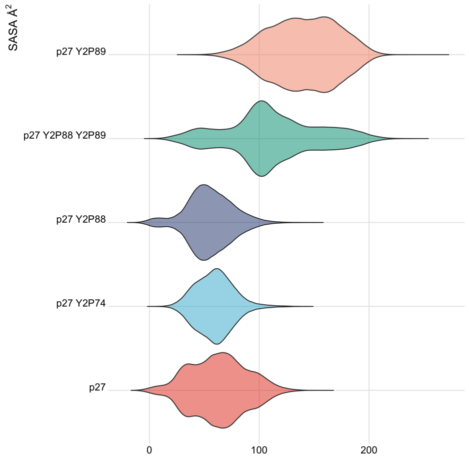


(b)

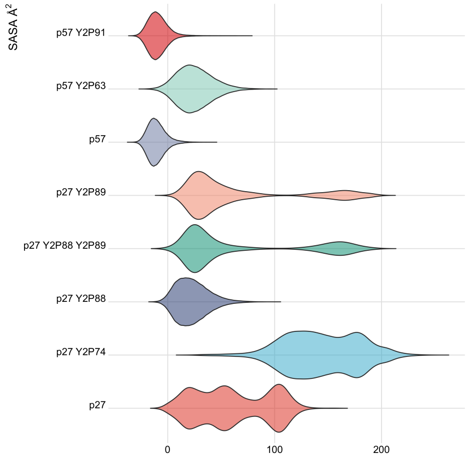


(c)

Supplementary Figure 5: Density of SASA of residue (a) p27$^{Y88}$, p57$^{Y91}$, p21$^{Y77}$, (b) p27$^{Y89}$ and (c) p27$^{Y74}$ and p57$^{Y63}$, taken from three independent cMD simulations for each model.

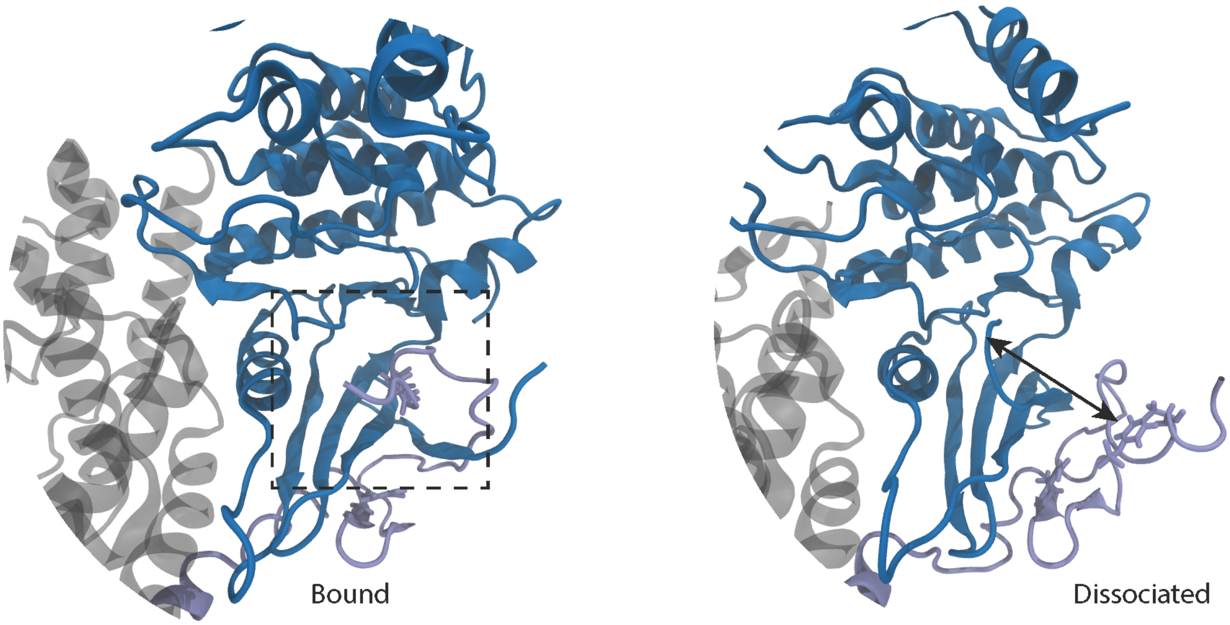


Supplementary Figure 6: The starting conformation of p57 bound to CyclinA/Cdk2, where we have highlighted the $p57^{Y91}$ residue tightly bound to the Cdk2 active site (left), and the final conformation from simulation highlighting the dissociation of $p57^{Y91}$ from Cdk2 and apparent reactivation of Cdk2.

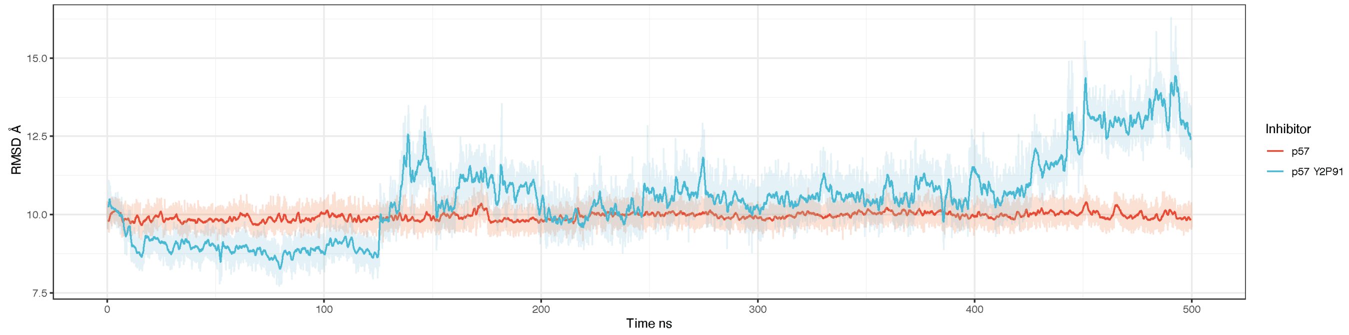


(a)

Supplementary Figure 7: Centre-of-mass distance between the $3_{10}$ $\alpha$-helicies and the active site pocket for p57 from cMD. The only complex displaying dissociation of the helix from classical molecular dynamics.

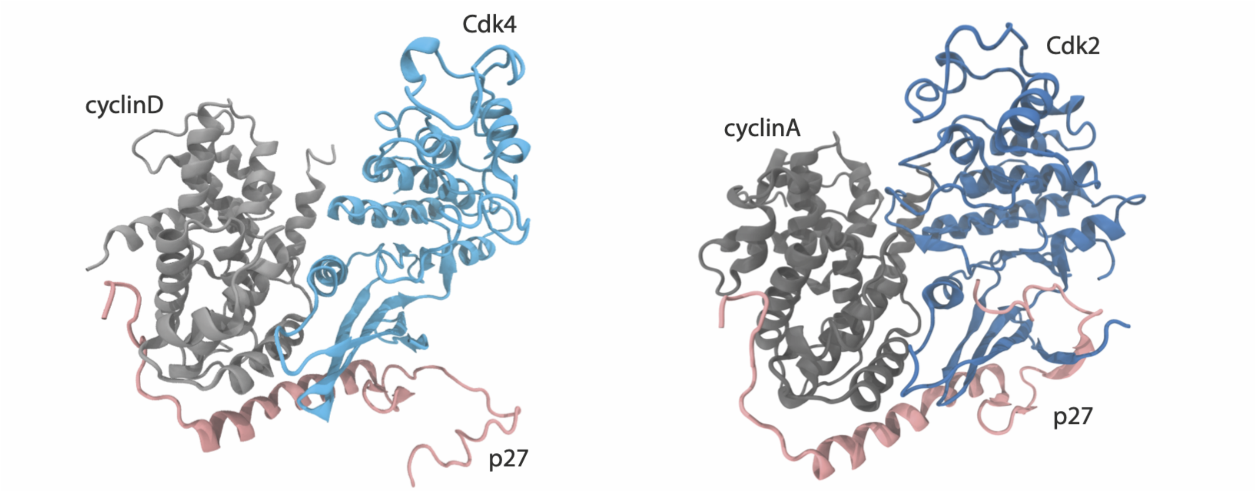


Supplementary Figure 8: A side-by-side comparison of the p27 bound Cdk2/CyclinA and Cdk4/CyclinD complexes.

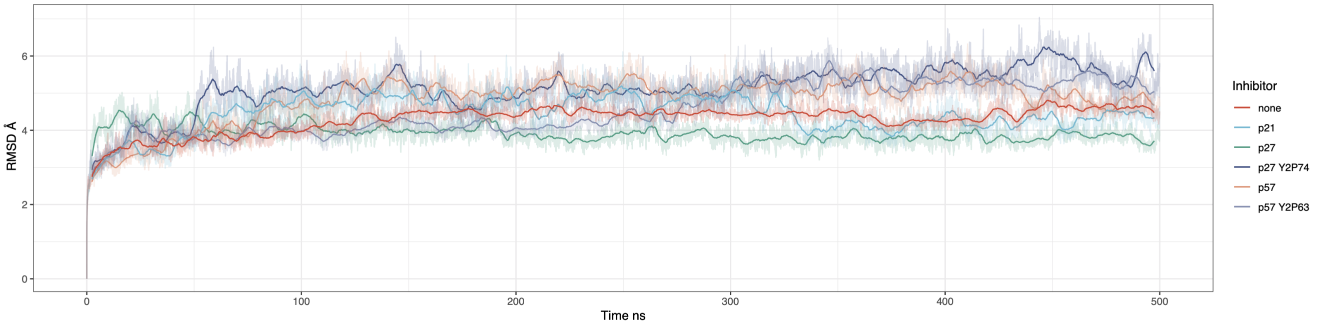


(a) Whole system

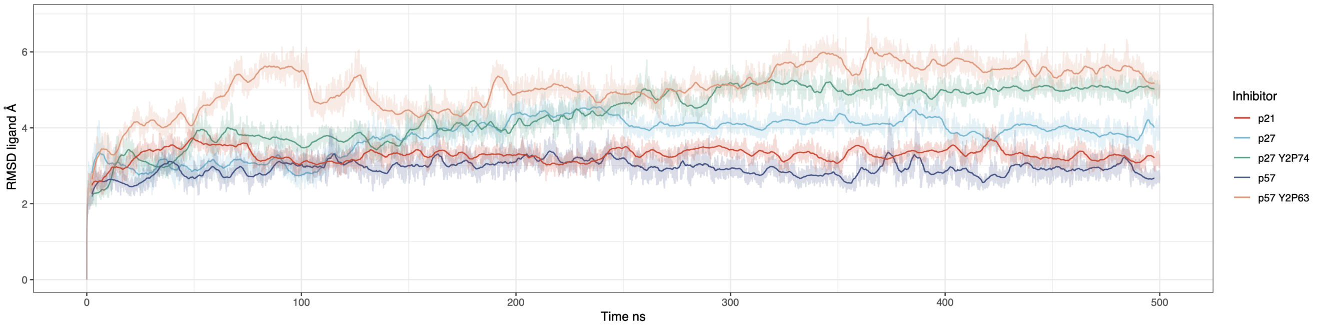


(b) Inhibitor only

Supplementary Figure 9: Root mean squared deviation from the crystal structure after minimisation and heat-up for each model of (a) the full complex and (b) the inhibitor. The p27 Y2P74 model displays greater deviation from it's starting structures than other models.

Supplementary Table 1: Simulation Details

| Model | Inhibitor | Cdk | Cyclin | cMD | aMD |
|---|---|---|---|---|---|
| 1 | none | Cdk2 | CyclinA | 500 ns | 500 ns |
| 2 | p27 | Cdk2 T2P13 | CyclinA | 500 ns | 500 ns |
| 3 | p27 | Cdk2 Y2P14 | CyclinA | 500 ns | 500 ns |
| 4 | p27 | Cdk2 T2P13/Y2P14 | CyclinA | 500 ns | 500 ns |
| 5 | p27 | Cdk2 | CyclinA | 1500 ns | 500 ns |
| 6 | p27 Y2P88 | Cdk2 | CyclinA | 1500 ns | 500 ns |
| 7 | p27 Y2P89 | Cdk2 | CyclinA | 1500 ns | 500 ns |
| 8 | p27 Y2P88 Y2P89 | Cdk2 | CyclinA | 1500 ns | 500 ns |
| 9 | p27 Y2P74 | Cdk2 | CyclinA | 1500 ns | 500 ns |
| 10 | p21 | Cdk2 | CyclinA | 1500 ns | 500 ns |
| 11 | p21 Y2P77 | Cdk2 | CyclinA | 1500 ns | 500 ns |
| 12 | p57 | Cdk2 | CyclinA | 1500 ns | 500 ns |
| 13 | p57 Y2P91 | Cdk2 | CyclinA | 1500 ns | 500 ns |
| 14 | p57 Y2P63 | Cdk2 | CyclinA | 1500 ns | 500 ns |
| 15 | none | Cdk4 | CyclinD | 500 ns | 500 ns |
| 16 | p27 | Cdk4 | CyclinD | 500 ns | 500 ns |
| 17 | p27 Y2P74 | Cdk4 | CyclinD | 500 ns | 500 ns |
| 18 | p21 | Cdk4 | CyclinD | 500 ns | 500 ns |
| 19 | p57 | Cdk4 | CyclinD | 500 ns | 500 ns |
| 20 | p57 Y2P63 | Cdk4 | CyclinD | 500 ns | 500 ns |

### Supplementary Table 2: CyclinA/Cdk2/p27 Intermolecular hydrogen bonds

| Acceptor | Donor | Donor H | Fraction | Avg Dist | Avg Ang |
|---|---|---|---|---|---|
| Cdk2 GLU81 O | p27 TYR88 HH | p27 TYR88 OH | 0.9277 | 2.7255 | 156.7414 |
| Cdk2 LYS20 O | p27 GLN77 H | p27 GLN77 N | 0.8497 | 2.8389 | 159.9591 |
| p27 ARG30 O | CycA GLN254 HE21 | CycA GLN254 NE2 | 0.7614 | 2.8550 | 163.6496 |
| p27 GLN77 O | Cdk2 LYS20 H | Cdk2 LYS20 N | 0.6941 | 2.8750 | 164.4870 |
| Cdk2 VAL18 O | p27 VAL79 H | p27 VAL79 N | 0.6316 | 2.8766 | 162.2807 |
| CycA THR285 OG1 | p27 ASN31 HD22 | p27 ASN31 ND2 | 0.5990 | 2.8777 | 163.1120 |
| p27 VAL79 O | Cdk2 VAL18 H | Cdk2 VAL18 N | 0.5775 | 2.8694 | 160.0881 |
| CycA GLU268 O | Cdk2 ARG150 HH12 | Cdk2 ARG150 NH1 | 0.5554 | 2.8259 | 155.3670 |
| CycA GLU269 O | Cdk2 ARG159 HH22 | Cdk2 ARG150 NH2 | 0.5444 | 2.8241 | 158.6409 |
| p27 ALA28 O | CycA TRP217 HE1 | CycA TRP217 NE1 | 0.5438 | 2.8429 | 152.3201 |

### Supplementary Table 3: CyclinA/Cdk2/p57 Intermolecular hydrogen bonds

| Acceptor | Donor | Donor H | Fraction | Avg Dist | Avg Ang |
|---|---|---|---|---|---|
| Cdk2 GLU81 O | p57 TYR91 HH | p57 TYR91 OH | 0.9252 | 2.7349 | 157.1118 |
| Cdk2 LYS20 O | p57 THR80 H | p57 THR80 N | 0.7832 | 2.8444 | 159.7401 |
| p57 ARG31 O | CycA GLN254 HE21 | CycA GLN254 NE2 | 0.7393 | 2.8585 | 163.2721 |
| p57 VAL82 O | Cdk2 VAL18 H | Cdk2 VAL18 N | 0.7028 | 2.8656 | 161.7094 |
| p57 THR80 O | Cdk2 LYS20 H | Cdk2 LYS20 N | 0.6906 | 2.8719 | 164.6074 |
| Cdk2 VAL18 O | p57 VAL82 H | p57 VAL82 N | 0.6027 | 2.8825 | 160.8764 |
| CycA GLU268 OE2 | Cdk2 ARG150 HH12 | Cdk2 ARG150 NH1 | 0.5722 | 2.8099 | 157.1202 |
| p57 ALA29 O | CycA TRP217 HE1 | CycA TRP217 NE1 | 0.5302 | 2.8593 | 156.4832 |

### Supplementary Table 4: CyclinA/Cdk2/p21 Intermolecular hydrogen bonds

| Acceptor | Donor | Donor H | Fraction | Avg Dist | Avg Ang |
|---|---|---|---|---|---|
| Cdk2 GLU81 O | p21 TYR77 HH | p21 TYR77 OH | 0.8877 | 2.7510 | 155.7238 |
| Cdk2 LYS20 O | p21 GLU66 H | p21 GLU667 N | 0.8584 | 2.8434 | 160.1049 |
| p21 VAL68 O | Cdk2 VAL18 H | Cdk2 VAL18 N | 0.7242 | 2.8612 | 161.6964 |
| p21 ARG19 O | CycA GLN254 HE21 | CycA GLN254 NE2 | 0.7212 | 2.8605 | 163.2194 |
| Cdk2 VAL18 O | p21 VAL68 H | p21 VAL68 N | 0.7042 | 2.8720 | 162.7763 |
| p21 GLU66 O | Cdk2 LYS20 H | Cdk2 LYS20 N | 0.6416 | 2.8808 | 165.1146 |
| p21 ALA17 O | CycA TRP217 HE1 | CycA TRP217 NE1 | 0.5339 | 2.8445 | 151.2931 |
| p21 ASP62 O | Cdk2 ASN23 HD22 | Cdk2 ASN23 ND2 | 0.5275 | 2.8400 | 153.8981 |

### Supplementary Table 5: CyclinA/Cdk2 Intermolecular hydrogen bonds

| Acceptor | Donor | Donor H | Fraction | Avg Dist | Avg Ang |
|---|---|---|---|---|---|
| CycA GLU269 O | Cdk2 ARG150 HH22 | Cdk2 ARG150 NH2 | 0.6919 | 2.8348 | 159.3739 |

### Supplementary Table 6: CyclinD/Cdk4/p27 Hbond details

| Acceptor | Donor | Donor H | Fraction | Avg Dist | Avg Ang |
|---|---|---|---|---|---|
| p27 GLN77 O | Cdk4 LYS24 H | Cdk4 LYS24 N | 0.8241 | 2.8484 | 162.4492 |
| CycD ALA153 O | Cdk4 ARG61 HH11 | Cdk4 ARG61 NH1 | 0.8058 | 2.8270 | 159.2312 |
| p27 ARG30 O | CycD GLN100 HE21 | CycD GLN100 NE2 | 0.7887 | 2.8483 | 163.3430 |
| Cdk4 LYS24 O | p27 GLN77 H | p27 GLN77 N | 0.7069 | 2.8672 | 162.2508 |
| CycD LYS149 O | Cdk4 ARG60 HE | Cdk4 ARG60 NE | 0.6619 | 2.8316 | 152.9677 |
| Cdk4 ASP128 OD2 | CycD ARG26 HE | CycD ARG26 NE | 0.6271 | 2.8293 | 160.7100 |
| Cdk4 ASP128 OD1 | CycD ARG26 HH21 | CycD ARG26 NH2 | 0.6240 | 2.8094 | 158.9971 |
| CycD GLU141 OE2 | Cdk4 LEU48 H | Cdk4 LEU48 N | 0.5566 | 2.8669 | 162.4664 |

### Supplementary Table 7: CyclinD/Cdk4/p57 Hbond details

| Acceptor | Donor | Donor H | Fraction | Avg Dist | Avg Ang |
|---|---|---|---|---|---|
| p57 THR80 O | Cdk4 LYS24 H | Cdk4 LYS24 N | 0.7715 | 2.8547 | 161.5388 |
| CycD LYS149 O | Cdk4 ARG60 HH11 | Cdk4 ARG60 NH1 | 0.7577 | 2.8222 | 158.0258 |
| p57 ARG31 O | CycD GLN100 HE21 | CycD GLN100 NE2 | 0.7568 | 2.8580 | 163.1346 |
| Cdk4 ASP128 OD2 | CycD ARG26 HH22 | CycD ARG26 NH2 | 0.6672 | 2.7841 | 161.6650 |
| Cdk4 ASP27 OD1 | p57 ARG76 HH22 | p57 ARG76 NH2 | 0.6490 | 2.7876 | 161.4593 |
| Cdk4 ASP128 OD1 | CycD ARG26 HH12 | CycD ARG26 NH1 | 0.6158 | 2.8135 | 161.4324 |
| Cdk4 LYS24 O | p57 THR80 H | p57 THR80 N | 0.6010 | 2.8724 | 160.2666 |
| Cdk4 ASP27 OD2 | p57 ARG76 HH12 | p57 ARG76 NH1 | 0.5057 | 2.8207 | 155.3517 |

### Supplementary Table 8: CyclinD/Cdk4/p21 Hbond details

| Acceptor | Donor | Donor H | Fraction | Avg Dist | Avg Ang |
|---|---|---|---|---|---|
| p21 GLU66 O | Cdk4 LYS24 H | Cdk4 LYS24 N | 0.8738 | 2.8409 | 162.3939 |
| p21 ARG19 O | CycD GLN100 HE21 | CycD GLN100 NE2 | 0.8019 | 2.8466 | 163.5126 |
| Cdk4 LYS24 O | p21 GLU66 H | p21 GLU66 N | 0.6739 | 2.8733 | 161.0108 |
| CycD LYS149 O | Cdk4 ARG60 HE | Cdk4 ARG60 NE | 0.5691 | 2.8463 | 155.4499 |
| Cdk4 ASP75 OD2 | p21 TRP49 HE1 | p21 TRP49 NE1 | 0.5166 | 2.8326 | 163.6698 |

### Supplementary Table 9: CyclinD/Cdk4 Hbond details

| Acceptor | Donor | Donor H | Fraction | Avg Dist | Avg Ang |
|---|---|---|---|---|---|
| Cdk4 ASP128 OD2 | CycD ARG26 HH22 | CycD ARG26 NH2 | 0.9339 | 2.7818 | 160.7384 |
| Cdk4 ASP128 OD1 | CycD ARG26 HH12 | CycD ARG26 NH1 | 0.8858 | 2.8056 | 163.3988 |
| CycD LYS149 O | Cdk4 ARG60 HE | Cdk4 ARG60 NE | 0.6789 | 2.8408 | 156.3797 |
| CycD GLU141 OE1 | Cdk4 LEU48 H | Cdk4 LEU48 N | 0.6267 | 2.8530 | 160.3516 |
